# Peptidoglycan-Chi3l1 interaction shapes gut microbiota in intestinal mucus layer

**DOI:** 10.1101/2023.10.03.560754

**Authors:** Yan Chen, Ruizhi Yang, Bin Qi, Zhao Shan

## Abstract

The balanced gut microbiota in intestinal mucus layer plays an instrumental role in the health of the host. However, the mechanisms by which the host regulates microbial communities in the mucus layer remain largely unknown. Here, we discovered that the host regulates bacterial colonization in the gut mucus layer by producing a protein called Chitinase 3-like protein 1 (Chi3l1). Intestinal epithelial cells are stimulated by the gut microbiota to express Chi3l1. Once expressed, Chi3l1 is secreted into the mucus layer where it interacts with the gut microbiota, specifically through a component of bacterial cell walls called peptidoglycan. This interaction between Chi3l1 and bacteria is beneficial for the colonization of bacteria in the mucus, particularly for gram-positive bacteria like *Lactobacillus*. Moreover, a deficiency of Chi3l1 leads to an imbalance in the gut microbiota, which exacerbates colitis induced by dextran sodium sulfate (DSS). By performing fecal microbiota transplantation from Villin-cre mice or replenishing *Lactobacillus* in IEC^ΔChil1^ mice, we were able to restore their colitis to the same level as that of Villin-cre mice. In summary, this study shows a “scaffold model” for microbiota homeostasis by interaction between intestinal Chi3l1 and bacteria cell wall interaction, and it also highlights that an unbalanced gut microbiota in the intestinal mucus contributes to the development of colitis.

## Introduction

Intestinal homeostasis is crucial for maintaining human health[1]. Alterations in gut microbiota composition have been linked to various diseases including cancer, obesity, and neurological disorders[2–6]. Dysbiosis, which refers to an imbalance in gut microbiota, is characterized by decreased microbial diversity, the presence of harmful microbes, or absence of beneficial ones [7]. Therefore, understanding the factors that influence gut microbiota is a fundamental goal in microbiome research[8]. Growing evidence suggests that colonization of the gut mucus layer can affect the susceptibility and progression of intestinal diseases like inflammatory bowel disease (IBD), irritable bowel syndrome, and celiac disease[9]. Inflammatory bowel diseases, such as Crohn’s disease (CD) and Ulcerative Colitis (UC), are characterized by chronic inflammation of the intestinal mucosa. Although the cause of the inflammatory bowel disease is unclear, <u>mouse</u> <u>models</u> lacking the key components of the mucus are predisposed to colitis, accompanied by dysbiosis in mucosa[10, 11], which is in accordance with increased epithelial-adherent microbial communities in biopsies from patients with IBD [12, 13]. Furthermore, there were significant differences in the gut microbiota of CD patients compared with healthy controls, and these differences were only present in mucus samples (not stool samples), suggesting that bacteria in the mucus layer may be more important for the development of IBD[14]. Donaldson et al. discovered that the intestinal flora utilizes host immunoglobulin A (IgA) for mucus colonization, indicating that the host may secrete certain factors to maintain intestinal flora homeostasis in the mucus[15]. However, the mechanisms regulating gut microbiota colonization in the mucus layer remain largely unknown. We aim to investigate the regulation of microbial communities in gut mucus and its implications in intestinal diseases.

Chitinase 3-like protein 1 (Chi3l1, also known as YKL-40 in humans) is a secreted protein that belongs to the glycosylhydrolase 18 family[16]. Despite its name, Chi3l1 can bind to chitin but does not have chitinase activity[17]. In our investigation, we noticed that chitin and peptidoglycan, a major component of bacterial cell walls, have similar structures [18]. Based on this information, we speculate that Chi3l1 might also interact with peptidoglycan and, therefore, interact with bacteria. Interestingly, Chi3l1 is expressed in intestinal epithelial cells and the lamina propria. We hypothesize that Chi3l1 may be secreted by intestinal epithelial cells and regulate the gut microbiota through its interaction with peptidoglycan in the mucous layer. In our study, we discovered that gut microbiota induce the expression of Chi3l1 in epithelial cells. Once expressed, Chi3l1 is secreted into the mucosa and interacts with bacteria, particularly with the bacterial cell wall component peptidoglycan. This interaction promotes bacterial colonization, especially of beneficial bacteria such as *Lactobacillus*. As a result, mice with higher levels of Chi3l1 are less susceptible to colitis.

## Results

### Intestinal epithelial cells express Chi3l1 induced by gut microbiota

The gut microbiota’s composition is shaped by host factors, including IgA [15], RegIIIγ [19], and TLR-5 [18]. Yet, specific host factors which maintain the homeostasis of the microbiota remain largely undefined. Drawing on the theory of co-evolution between the host and microbiota [20], we propose that host factors, which are induced by bacteria in the gut, could play a pivotal role in regulating bacterial colonization and composition. In a previous study, it was observed that both live *E.coli* and heat-killed *E.coli* treatment resulted in a significant increase in the expression of the gene encoding Chi3l1 in human intestinal organoids[21]. To verify this finding, we conducted immunohistochemical staining on intestinal tissue sections of germ-free and specific pathogen free (SPF) C57BL/6J wildtype mice. We observed a substantial increase of Chi3l1 expression in SPF mice compared to germ-free mice (Fig. 1A). The intestinal epithelium comprises various cell types, including intestinal cells, goblet cells, endoentercrine cells, Tuft cells, Paneth cells, M cells, and others [21]. To identify the cellular sources of Chi3l1, we performed co-staining with markers for specific cell types, including Chromogranin A (ChgA) for enteroendocrine cells, Ulex Europaeus Agglutinin I (UEA-1) for goblet cells and Paneth cells, and Double cortin-like kinase 1 (DCLK1) for tuft cells. Our results revealed that Chi3l1 is primarily expressed in enteroendocrine cells in the ileum and goblet or Paneth cells in the colon (Fig. 1B). However, Chi3l1 expression was not observed in tuft cells (Supplementary Figure 1).

**Figure 1.**
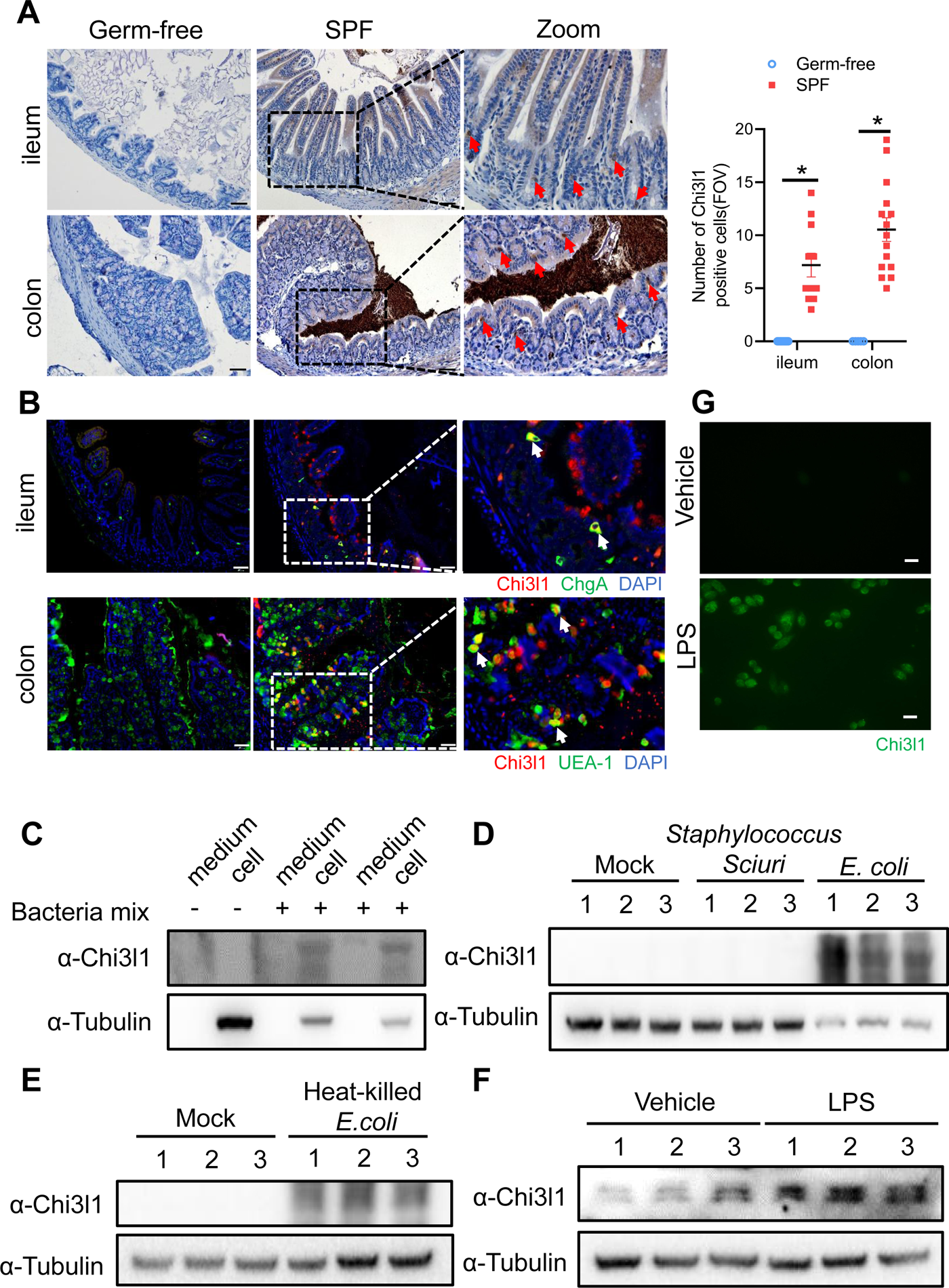
Intestinal epithelial cells express Chi3l1 induced by gut microbiota. **(A)** IHC staining to detect Chi3l1 in both ileum and colon from germ-free and wildtype mice. Ctrl (wildtype mice without application of first antibody), WT (wildtype C57B/6J mice). Red arrows indicate Chi3l1-expressing cells. Scale bar, 50μm (Ctrl, Germ-free, WT) and 20μm(zoom). <u>The number of Chi3l1-positive cells in each field of view (FOV) was analyzed.</u> **(B)** Ileum and colon were collected from wildtype mice and stained with ChgA (green), Chi3l1(red), and nuclear DAPI (blue) in ileum and UEA-1 (green), Chi3l1(red), and nuclear DAPI (blue) in colon. Scale bar, 20μm. Ctrl (without application of first antibody), WT (wildtype C57B/6J mice). **(C)** Western blot to detect Chi3l1 protein expression in DLD-1 cells after bacteria mix infection for 12 hour. Bacteria mix are total bacteria extracted from feces of wildtype mice. **(D)** Western blot to detect Chi3l1 protein expression in DLD-1 cells after *Staphylococcus Sciuri* and *E. coli* infection for 12 hour*. Staphylococcus Sciuri* and *E. coli* are isolated from bacteria mix and verified by 16S rRNA sequencing. Three independent experimental results are showed. **(E)** Western blot to detect Chi3l1 protein expression in DLD-1 cells after treatment with heat-killed *E. coli* for 12 hour. Three independent experimental results are showed. **(F)** Western blot to detect Chi3l1 protein expression in DLD-1 cells after treatment with 100 pg/ml LPS for 12 hour. Three independent experimental results are showed. **(G)** Immunofluorescence to detect Chi3l1 protein expression in DLD-1 cells after treatment with 100pg/ml LPS for 12 hour. Scale bar, 20μm. <u>The presence of cells in the</u> <u>untreated sample are annotated using white dashed lines based on the overexposure.</u> All data above represent at least three independent experiments. Representative images are shown in A,B, n=3-4 mice/group.

Furthermore, we isolated total bacteria from wildtype mouse feces and treated DLD-1 cells (a colorectal adenocarcinoma cell line) with the bacterial extract for 12 hours. We found that the bacterial extract directly induced Chi3l1 expression in DLD-1 cells (Fig. 1C). To examine whether the induction of Chi3l1 expression require a specific bacteria strain, we further identified the bacterial extract using 16S rRNA sequencing. Our results revealed that *E.coli* specifically stimulated Chi3l1 expression in DLD-1 cells, while *Staphylococcus Sciuri* did not have the same effect (Fig. 1D). <u>Although our data are limited to these two bacterial strains, it suggest that not all</u> <u>bacteria can induce the expression of Chi3l1.</u> Next, we wondered what component of bacteria can induce Chi3l1 expression. We tried heat-killed *E.coli*, which maintains bacterial cell wall integrity. We found that treatment of DLD-1 cells with heat-killed *E.coli* also led to an induction of Chi3l1 expression (Fig. 1E). These findings suggest that the induction of Chi3l1 expression does not necessarily require live bacteria and that bacterial components alone are sufficient to induce this response.

<u>Given that *E. coli* is Gram-negative and *Staphylococcus sciuri* is Gram-positive, we hypothesized that the difference in their ability to induce Chi3l1 expression might be due to variations between Gram-negative and Gram-positive bacteria, such as the presence of lipopolysaccharides (LPS). To test this hypothesis, we used LPS to induce Chi3l1 expression. Consistent with our hypothesis, LPS successfully induced Chi3l1 expression (Fig. 1F&G)</u>. Collectively, these findings provide evidence that the gut microbiota can induce Chi3l1 expression in intestinal epithelial cells. Collectively, these findings provide evidence that the gut microbiota can induce Chi3l1 expression in intestinal epithelial cells.

### Chi3l1 interact with bacteria via peptidoglycan

Chi3l1 belongs to a group of proteins called non-enzymatic chitinase-like proteins, which are known to bind chitin. Chitin is a polysaccharide present in the exoskeleton of arthropods and the cell walls of fungi[22]. By comparing the structure of chitin with that of peptidoglycan (PGN), a component of bacterial cell walls, we observed that they have similar structures (Fig. 2A). Based on this observation, we hypothesized that Chi3l1 may interact with gut bacteria through PGN. To test our hypothesis, we conducted co-incubation experiments where we mixed recombinant mouse Chi3l1 (rmChi3l1) with either Gram-positive or Gram-negative bacteria and then precipitated the bacteria through centrifugation. We found that rmChi3l1 was present in the pellet obtained from both Gram-positive and Gram-negative bacteria (Fig. 2B), suggesting that Chi3l1 can directly interact with bacteria.

**Figure 2.**
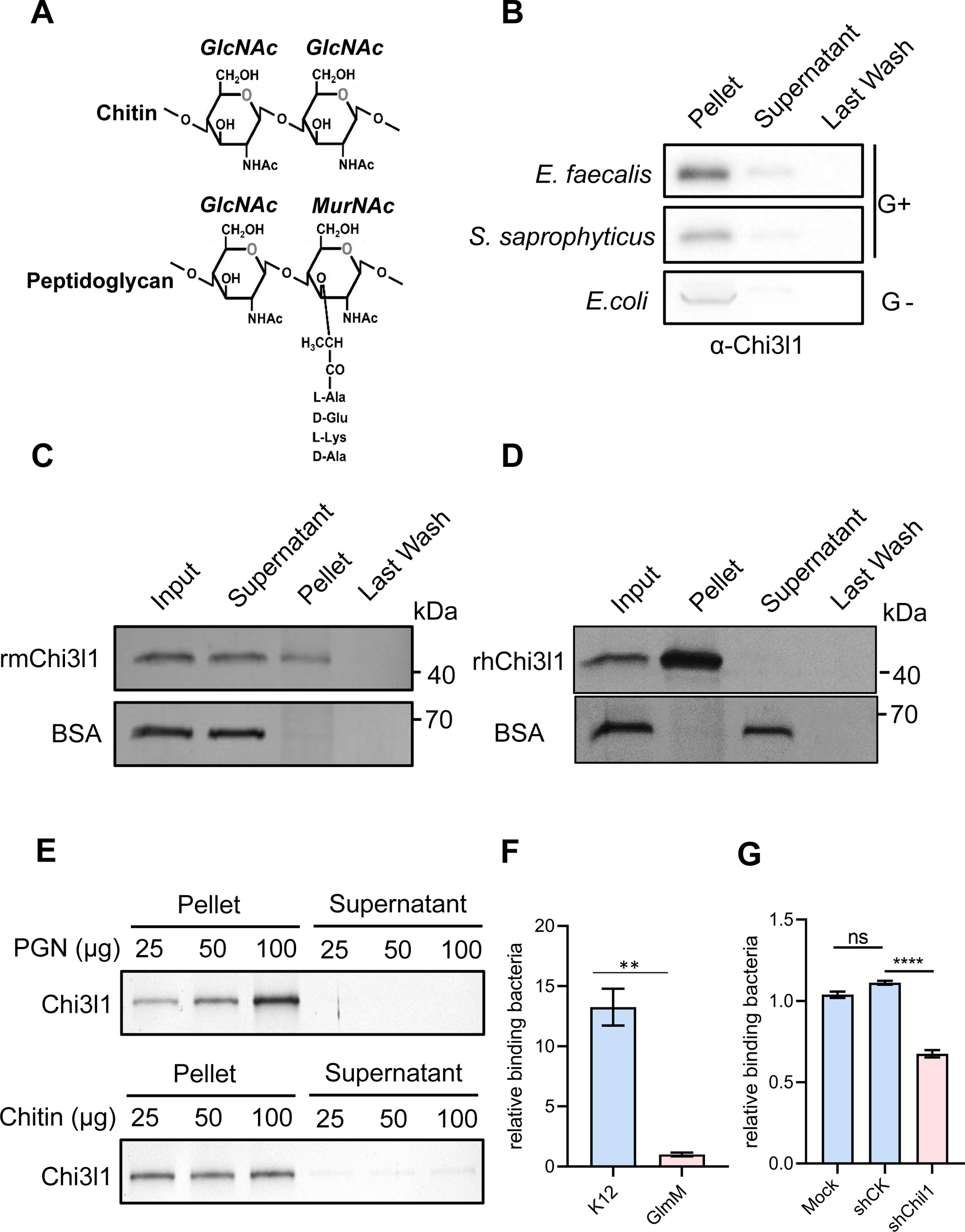
Chi3l1 interact with bacteria via peptidoglycan. **(A)** Structural comparison between Chitin and peptidoglycan.<u>Both chitin and peptidoglycan contain N-acetylglucosamine (GlcNAc) and have β-1,4-glycosidic bonds in their structures. However, Chitin is purely a polysaccharide, while peptidoglycan includes a peptide component that forms cross-links between chains</u>. [30] **(B)** Gram-positive bacteria (*E.faecalis*, *S.saprophyticus*) and Gram-negative bacteria (*E.coli*) were incubated with 1μg of recombinant mouse Chi3l1 protein(rmChi3l1), respectively. Proteins bound to indicated bacteria were precipitated by centrifugation. Western blot was used to detect rmChi3l1 in Pellet, Supernatant (unbound proteins) and Last Wash (last wash unbound proteins). **(C)** Insoluble Peptidoglycan (PGN) were incubated with either recombinant mouse Chi3l1 protein (rmChi3l1) or Bovine serum albumin (BSA). Proteins bound to PGN were precipitated by centrifugation. Silver staining was used to detect rmChi3l1 in Input, Supernatant (unbound proteins), Pellet and Last Wash (last wash unbound proteins). **(D)** Insoluble Peptidoglycan (PGN) were incubated with recombinant human Chi3l1 protein (rhChi3l1). Proteins bound to PGN were precipitated by centrifugation. Silver staining was used to detect rhChi3l1 in Input, Supernatant (unbound proteins), Pellet and Last Wash (last wash unbound proteins). All data above represent at least three independent experiments. **<u>(E)</u>** <u>Insoluble</u> <u>PGN or chitin was incubated with rmChi3l1. Chi3l1 bound to PGN (upper panel) and chitin</u> <u>(lower panel) was precipitated and detected by silver staining. The supernatant represents the last</u> <u>wash, and the pellet contains proteins precipitated by either PGN or chitin. **(F)** Relative DLD-1</u> <u>bacterial binding preference after treatment with K12 or GlmM, a PGN synthesis-deficient mutant.</u> <u>Colony-forming units (CFU) were counted, and GlmM CFU were normalized to 1. **(G)** Relative</u> <u>K12 bacterial adhesion preference after DLD-1 cells were transfected without (Mock), or with</u> <u>scramble shRNA (shCK), or with shChil1. CFU were counted, and the Mock group were</u> <u>normalized to 1.</u>

To further investigate the interaction between Chi3l1 and PGN, we also co-incubated PGN with rmChi3l1 and precipitated the PGN through centrifugation. PGN is an insoluble substance and hence can be precipitated by centrifugation. Consistent with our previous results, we observed that rmChi3l1 was present in the pellet obtained from PGN but not in the pellet obtained from bovine serum albumin (BSA), which served as a negative control (Fig. 2C). Furthermore, we also examined the interaction between PGN and recombinant human Chi3l1 (rhChi3l1) and obtained similar results (Fig. 2D). These findings indicate that Chi3l1 interacts with bacteria through PGN.

<u>To better characterize the binding between Chi3l1 and peptidoglycan (PGN), we compared the binding affinities of Chi3l1 to both PGN and chitin. We incubated chitin and PGN with rmChi3l1 in increasing doses (25, 50, 100 μg) and detected the precipitated rmChi3l1 by either chitin or PGN. Our results indicate that Chi3l1 interacts with PGN in a dose-dependent manner (Figure 2E). In contrast, the binding between Chi3l1 and chitin did not exhibit dose dependency (Figure 2E). These findings suggest a specific and distinct binding mechanism for Chi3l1 with PGN compared to chitin</u>.

<u>To investigate whether the Chi3l1-PGN interaction could facilitate the colonization of gram-positive bacteria, we conducted adhesion experiments using DLD-1 cells and bacteria. We employed a GlmM mutant (PGN synthesis-deficient) and K12 bacteria (a wild-type E. coli strain used as a control) to assess their adhesion capabilities. The results showed that the adhesion ability of the GlmM mutant to cells significantly decreased (Figure 2F). Additionally, after knocking down Chi3l1 in DLD-1 cells (knockdown efficiency over 50%), we observed a decrease in bacterial adhesion (Figure 2G). These findings suggest that the Chi3l1-PGN interaction plays a crucial role in bacterial adhesion</u>.

### Intestinal bacteria are disordered in IEC^ΔChil1^ mice, especially Gram-positive bacteria

To gain initial insights into how the expression of Chi3l1 in intestinal epithelial cells (IECs) affects the colonization of gut microbiota, we created mice with a specific deficiency of Chi3l1 in IECs (referred to as IEC^ΔChil1^ mice) (Supplementary Figure 2). We then conducted bacterial 16S rRNA sequencing analysis of the colon contents of both Villin-cre and IEC^ΔChil1^ littermates. Our analysis of alpha diversity revealed that the bacterial population was relatively lower in IEC^ΔChil1^ littermates compared to Villin-cre littermates (Figure 3A). This finding was further confirmed by conducting universal bacterial 16S rRNA qPCR analysis of the feces and ileum contents of IEC^ΔChil1^ and Villin-cre littermates, which also showed lower bacterial enrichment in IEC^ΔChil1^ mice (Figure 3B). Furthermore, principal component analysis demonstrated significant differences in bacterial diversity between Villin-cre and IEC^ΔChil1^ littermates (Figure 3C).

**Figure 3.**
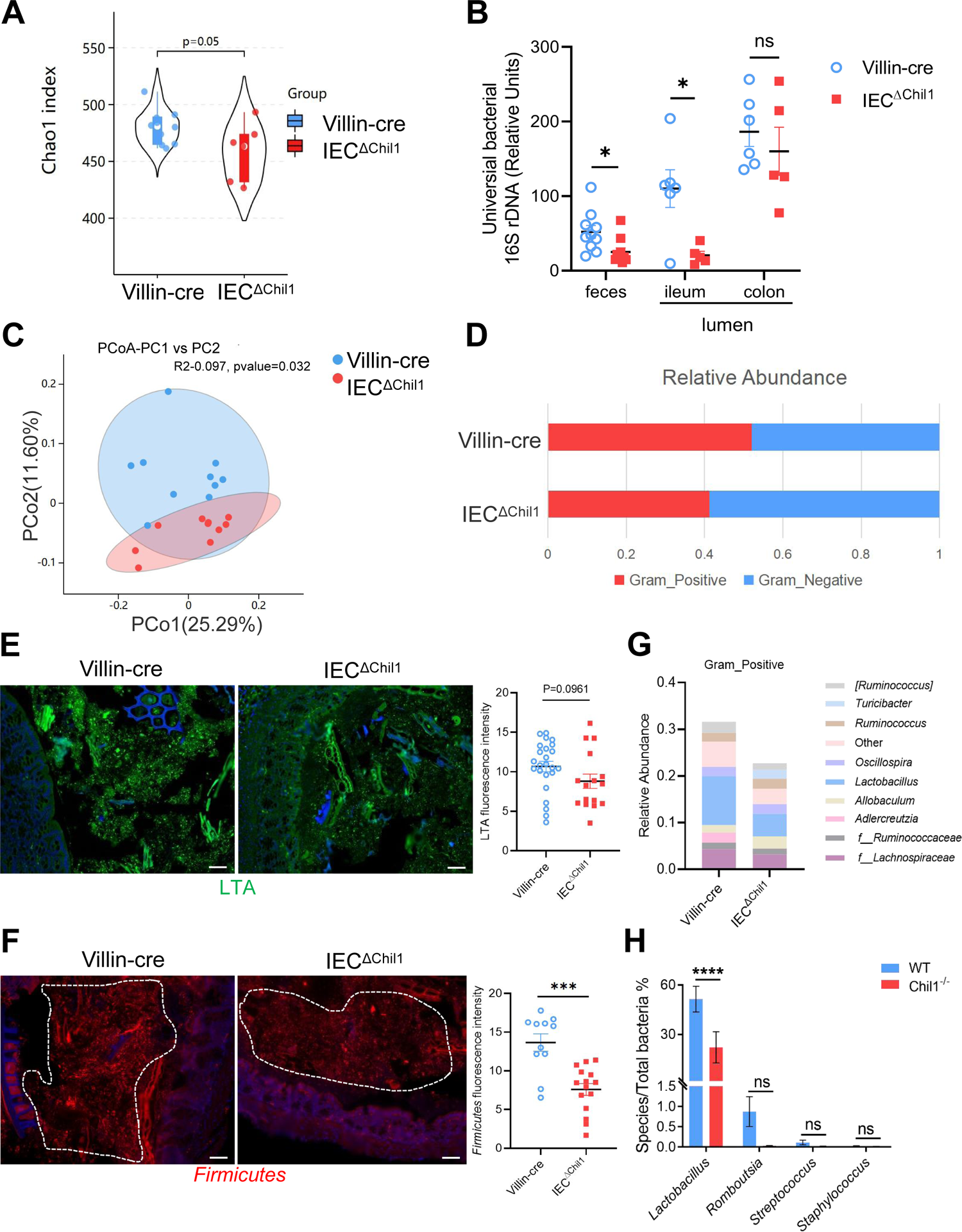
Intestinal bacteria are disordered in IEC^ΔChil1^ mice, especially Gram-positive bacteria. **(A, C, D, G)** Female Villin-cre and IEC^ΔChil1^ littermates continue to cage together after weaning for 8 weeks. Microbial communities in feces and intestinal lumen were characterized by 16S rRNA sequencing. n=7 or 10/group. **(A)** Alpha diversity analysis of colon contents between Villin-cre and IEC^ΔChil1^ littermates. **(B)** qPCR analysis of total bacteria in the feces and ileum, colon luminal microbial communities of Villin-cre and IEC^ΔChil1^ littermates. Values for each bacterial group are expressed relative to total 16S rRNA levels. Mean ± SEM is displayed. Two-tailed, unpaired student t-test was performed. *P<0.05, ns, not significant. n=5-10/group. **(C)** Principal Component Analysis of weighed UniFrac distances of 16S community profiles of Villin-cre and IEC^ΔChil1^ littermates feces (binary-jaccard). **(D)** Relative abundance of Gram-positive and Gram-negative bacteria in colon contents of Villin-cre and IEC^ΔChil1^ littermates are shown. **(E)** LTA (green) was detected by immunofluorescence in colon sections of Villin-cre and IEC^ΔChil1^ littermates. Nuclei were detected with DAPI. Scale bar, 50μm. <u>The average fluorescence intensity in each field of view (FOV) was analyzed.</u> **(F)** Fluorescence in situ hybridization (FISH) detection of gram-positive bacteria (red) in the colon of Villin-cre and IEC^ΔChil1^ littermates, nuclei were detected with DAPI (blue). Scale bars, 50um. The average fluorescence intensity in each field of view (FOV) was analyzed. **(G)** Relative abundance of Gram-positive bacteria genera in colon lumen of Villin-cre and IEC^ΔChil1^ littermates. **(H)** Female wildtype and Chil1^-/-^ littermates continue to cage together after weaning for 8 weeks. Microbial communities in feces were characterized by 16S rRNA sequencing. Mean ± SEM is displayed. Two-tailed, unpaired student t-test was performed. ****P<0.0001, ns, not significant. n=4 or 6/group. Representative images are shown in E, F, n= 3/4 mice/group.

When we examined the relative abundance of Gram-positive and Gram-negative bacteria between Villin-cre and IEC^ΔChil1^ littermates, we observed that Gram-positive bacteria were significantly reduced in IEC^ΔChil1^ mice, while there was no notable difference in Gram-negative bacteria (Figure 3D). This result was further validated by staining lipoteichoic acid (LTA), a component present on Gram-positive bacteria, which revealed a lower abundance of Gram-positive bacteria in IEC^ΔChil1^ compared to Villin-cre littermates (Figure 3E). Moreover, visualization of *Firmicutes* by bacteria FISH staining, a dominant group of Gram-positive bacteria in the gut, also showed reduced levels of *Firmicutes* in the colon lumen of IEC^ΔChil1^ mice compared to Villin-cre mice (Figure 3F). Analysis of the relative abundance of specific Gram-positive bacterial species demonstrated a significant reduction in *Lactobacillus* in IEC^ΔChil1^ mice compared to Villin-cre mice (Figure 3G). Similar results were observed in Chil1^-/-^ mice compared to wildtype mice (Figure 3H). <u>Consist</u> <u>with the 16S rRNA sequencing analysis data, qPCR results showed that *Turicibacter* was more</u> abundant in IEC^ΔChil1^ mice than Villin-cre mice (Supplementary Figure 3A). These findings suggest that Chi3l1 plays a role in regulating the colonization of Gram-positive bacteria, particularly *Lactobacillus*, in the murine gut.

### Chi3l1 promotes the colonization of Gram-positive bacteria in intestinal mucus

Chi3l1 was found in secretory cells like goblet cells and Paneth cells, suggesting that it may be secreted into the intestinal lumen (Fig. 1B). Immunohistochemistry staining of Chi3l1 in the colon revealed a large amount of Chi3l1 signals in the mucus layer (Fig. 4A). Immunofluorescence co-staining of Chi3l1 with UEA-1 in the colon yielded similar results (Fig. 4B). Furthermore, Chi3l1 was also detected in the ileum and colonic tissues and mucus layer (Fig. 4C&D). These findings indicate that mouse Chi3l1 is specifically expressed in intestinal secretory epithelial cells and secreted into the intestinal lumen. Since large amounts of Chi3l1 is secreted into the mucus and Chi3l1 interact with bacteria, we hypothesize that Chi3l1 may regulate the colonization of Gram-positive bacteria in the mucus layer. To test this hypothesis, we isolated bacterial DNA from the ileum and colon mucus of both wildtype and Chi11^-/-^ mice. Quantification of Gram-positive bacteria and *Lactobacillus* using qPCR revealed that both the ileum and colon mucus of Chi11^-/-^ mice had significantly lower levels of Gram-positive bacteria and *Lactobacillus* compared to that of wildtype mice (Fig. 4E&F).

**Figure 4.**
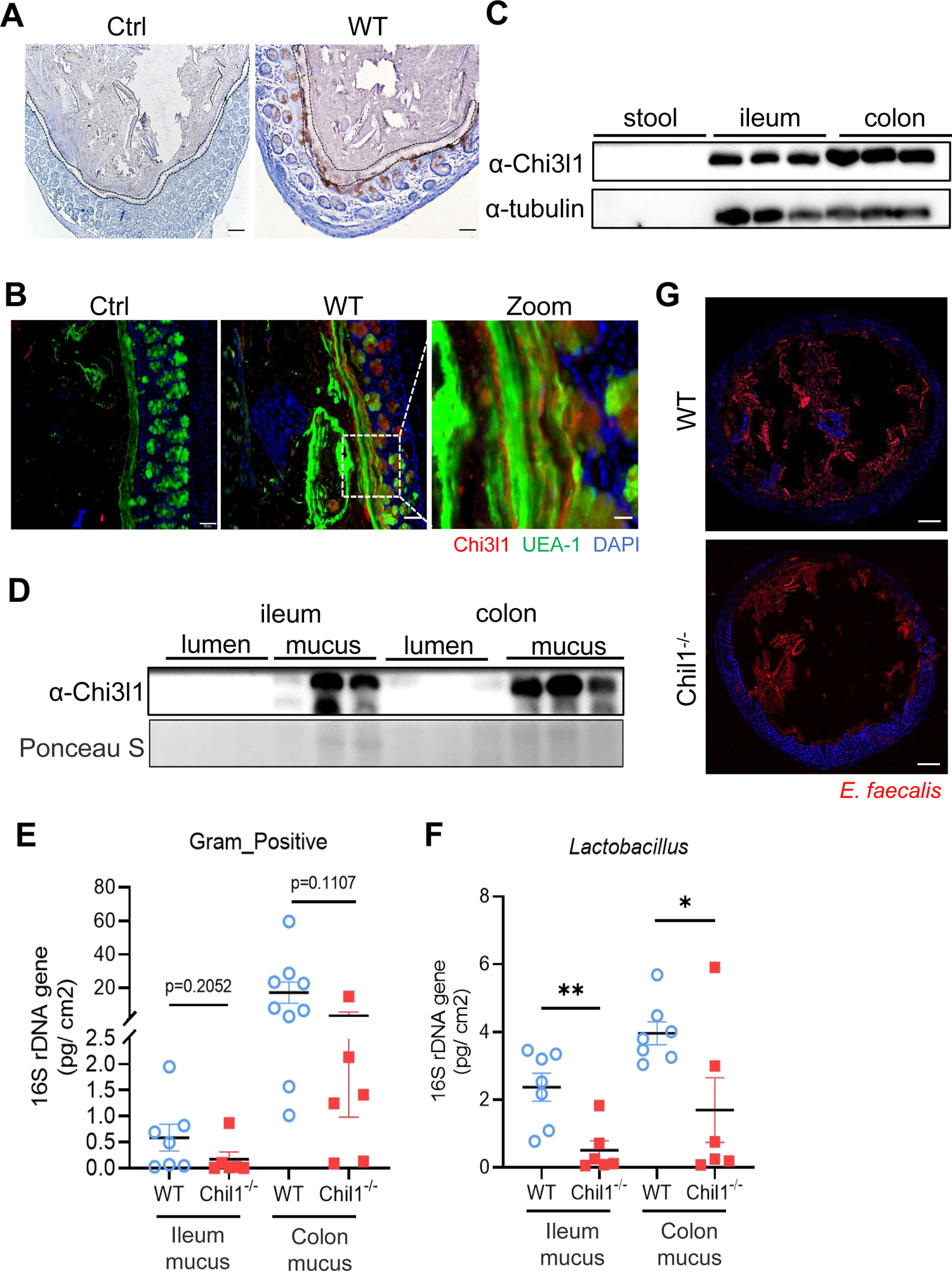
Chi3l1 promotes the colonization of Gram-positive bacteria in intestinal mucus layer. **(A)** IHC staining to detect Chi3l1 in colon mucus layer from wildtype mice. Ctrl (without application of ant-Chi3l1 antibody), WT (wildtype C57B/6J mice). Black dotted line outlines mucus layer. Scale bar, 50μm (Ctrl, WT). **(B)** Colons were collected from wildtype mice and stained with UEA-1 (green), Chi3l1(red), and nuclear DAPI (blue). Ctrl (without application of first antibody), WT (wildtype C57B/6J mice). Scale bars, 50um (Ctrl, WT) and 20μm(zoom). **(C)** Stool, ileum and colon tissues were collected from wildtype mice. Western blot was used to detect Chi3l1 expression in these samples. n=3 mice/sample. **(D)** Both luminal and mucus-associated proteins of either ileum or colon were extracted. Western blot was used to detect Chi3l1 expression in these samples. lumen (luminal proteins), mucus (mucus-associated proteins). n=3 mice/sample. **(E&F)** qPCR analysis of specific bacteria in the ileum and colon mucus microbial communities of wildtype and Chil1^-/-^ littermates. **(E)** qPCR analysis of Gram-positive bacteria is shown. **(F)** qPCR analysis of Gram-positive bacteria is shown. Values for each bacterial group are expressed relative to total 16S rRNA levels. WT (wildtype C57B/6J mice). Mean ± SEM is displayed. Two-tailed, unpaired student t-test was performed. P value is as indicated in E. *P<0.05, **P<0.01, n=6-9/group. **(G)** Rectal injection of both wildtype and Chi11^-/-^ mice with FDAA-labeled *E. faecalis* (a Gram-positive bacteria strain) for 4 hour. Colon sections were collected and colonization of *E.faecalis* was examined under microscope. Nuclei were stained with DAPI. Representative images are shown in A, B, G, n=3-4 mice/group.

To further validate these results, we labeled a Gram-positive bacteria strain, *E. faecalis*, with fluorescent D-amino acids (FDAA), which can metabolically label bacterial peptidoglycans[23]. We then performed rectal injection of both wildtype and Chi11^-/-^ mice with FDAA-labeled *E. faecalis*. The data demonstrated that Chi11^-/-^ mice had much lower colonization of *E. faecalis* compared to wildtype mice (Fig. 4G). <u>Besides Gram-positive bacteria, we also performed rectal</u> injection of both wildtype and Chi11^-/-^ mice with mCherry-OP50 (*a strain of E.coli that express mCherry*), we found Chi11^-/-^ mice had much higher colonization of *E. coli* compared to wildtype <u>mice (</u>Supplementary Figure 3B). Based on these findings, we conclude that Chi3l1 promotes the colonization of Gram-positive bacteria, particularly *Lactobacillus*, in the intestinal mucus. Additionally, we also observed that the deletion of Chi3l1 significantly reduced mucus layer thickness, which may be attributed to the disrupted colonization of Gram-positive bacteria in the intestinal mucus layer (Supplementary Figure 3 A&B).

### Disordered intestinal bacteria in IEC^ΔChil1^ mice contribute to colitis

From the above data, it is evident that Chi3l1 is secreted into the intestinal mucus to influence the colonization of Gram-positive bacteria, particularly *Lactobacillus*. We are now interested in understanding the role of Chi3l1-regulated microbiota in a pathological condition. We observed a significant increase in *Chil1* mRNA expression in the colon tissues of patients with either Crohn’s disease (CD) or Ulcerative colitis (UC) compared to normal tissues (Fig. 5A). To investigate further, we created a colitis mouse model by subjecting Villin-cre and IECΔChil1 mice to a 2% dextran sulfate sodium (DSS) diet for 7 days (Fig. 5B). The severity of colitis was assessed based on weight loss, colon length, and tissue damage. Without the DSS challenge, the colon length and structure were similar between Villin-cre and IEC^ΔChil1^ mice (Fig. 5D&E). However, upon DSS challenge, the IEC^ΔChil1^ mice showed significantly shorter colon length, faster body weight loss, and more severe inflammation compared to the Villin-cre mice (Fig. 5C-E).

**Figure 5.**
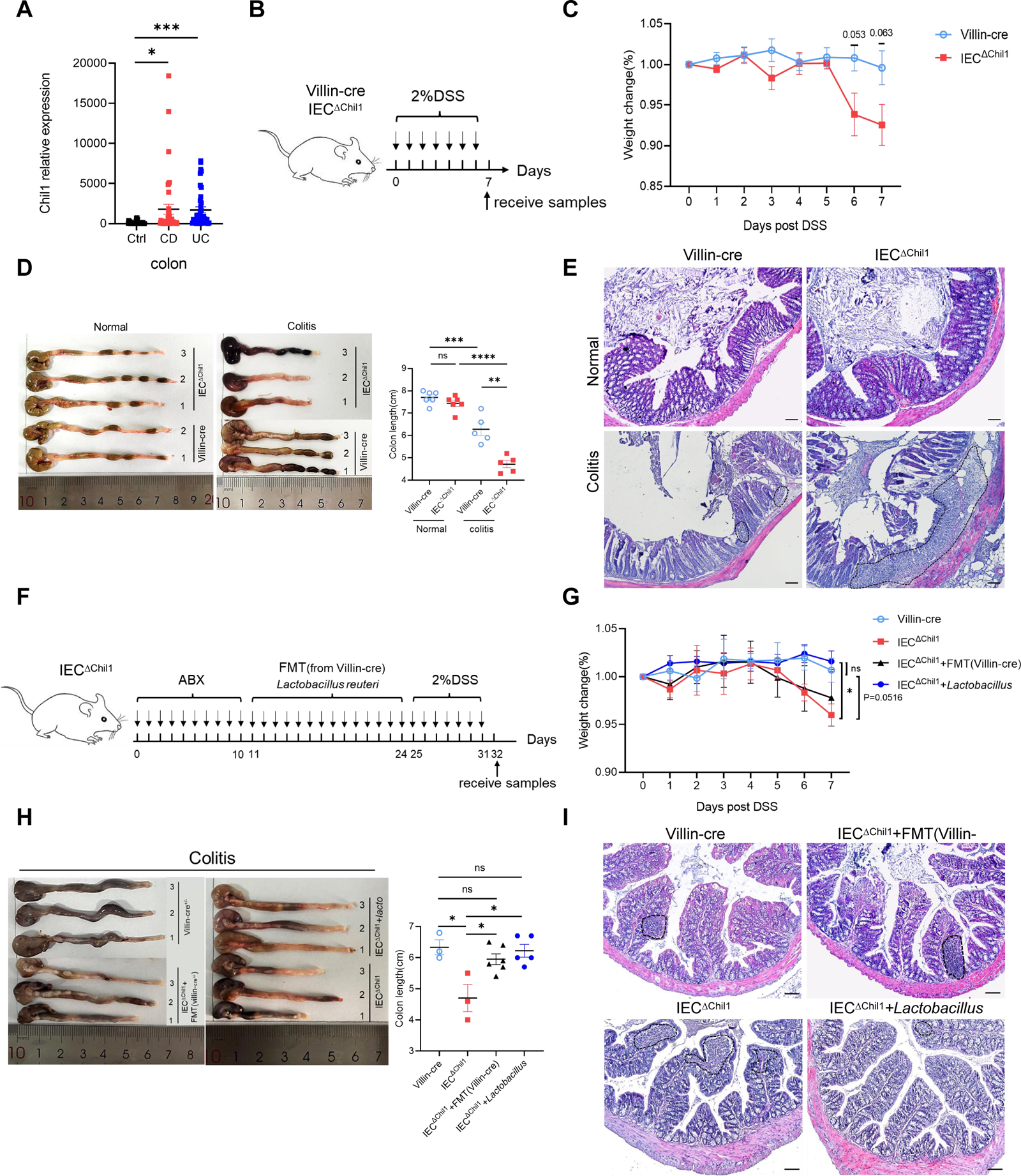
Disordered intestinal bacteria in IEC^ΔChil1^ mice contribute to IBD. **(A)** *Chil1* mRNA relative expression in colon tissues of patients without gut disease (controls, n=35) or with Crohn’s disease (CD, n=40), ulcerative colitis (UC, n=40), (GEO Datasets: SRP303290), Mann-Whitney test was performed. *P < 0.05; ***P < 0.001. **(B)** Schematic model of the experimental design. Both Villin-cre and IEC^ΔChil1^ littermates were fed with 2% DSS in drinking water to induce colitis. **(C)** Weight change of Villin-cre and IEC^ΔChil1^ littermates during DSS feeding. <u>Weight change</u> (%) = Current weight/Initial weight. P values are as indicated. **(D)** Representative colonic length from Normal and DSS-treated Villin-cre and IEC^ΔChil1^ littermates (left) and the statistics of colonic length (right) (**P < 0.01; ***P < 0.001; ****P < 0.0001, ns, not significant). **(E)** H&E staining of mice colon from Normal and DSS-treated Villin-cre and IEC^ΔChil1^ littermates. The inflamed areas are outlined by white dotted line, Scale bars=100um. **(F)** Schematic of the experimental design. First, antibiotics were used to eliminate gut microbiota for 10 days, and then either fecal microbiota from Villin-cre mice (FMT) or *Lactobacillus reuteri* were transplanted back *to* IEC^ΔChil1^ mice orally every day for 2 weeks. Finally, colitis mouse model was constructed by 2% DSS feeding in drinking water for another 7 days. **(G-I)** Villin-cre and IEC^ΔChil1^ were only fed with 2% DSS in drinking water for 7days. IEC^ΔChil1^+FMT(Villin-cre), and IEC^ΔChil1^+*Lactobacillus* were constructed as described in F. **(G)** <u>Weight change</u> of Villin-cre, IEC^ΔChil1^, IEC^ΔChil1^+FMT(Villin-cre), and IEC^ΔChil1^+*Lactobacillus* mice during DSS feeding. **(H)** Representative colonic length from Villin-cre, IEC^ΔChil1^, IEC^ΔChil1^+FMT(Villin-cre), and IEC^ΔChil1^+*Lactobacillus* mice (left) and the statistics of colonic length (right). Mean ± SEM is displayed in G, H. one-way ANOVA was performed. P value is as indicated. *P<0.05, ns, not significant, n=3-6/group. **(I)** H&E staining of mice colon from Villin-cre, IEC^ΔChil1^, IEC^ΔChil1^+FMT(Villin-cre), and IEC^ΔChil1^+*Lactobacillus* mice after DSS treatment. The inflamed area is outlined by black dotted line, Scale bars=100um. Representative images are shown in C, E, H, I, n=3-6 mice/group.

To rule out the effects of Chi3l1 on the host contributed to colitis, we pretreated the mice with antibiotics to eliminate gut microbiota before inducing colitis (Supplementary Figure 4A). The Universal bacterial 16S rRNA qPCR data indicated that the majority of gut microbiota were eliminated after antibiotics treatment (Supplementary Figure 4B). However, the IEC^ΔChil1^ mice exhibited a milder colitis phenotype, including slower body weight loss, longer colon length, and less inflammation compared to the Villin-cre mice (Supplementary Figure 4C-E). We believe that this could be due to the relationship between Chi3l1 and inflammation. Based on these findings, it is apparent that Chi3l1’s effects on gut microbiota play a more significant role in colitis.

To further elucidate the role of Chi3l1-regulated gut microbiota in colitis, we conducted fecal microbiota transplantation (FMT) and *Lactobacillus reuteri* transplantation experiments. We first eliminated gut microbiota through a 10-day course of antibiotics and then performed FMT from Villin-cre mice or administered oral gavage of *Lactobacillus reuteri* to IEC^ΔChil1^ mice for 2 weeks, followed by a 7-day period of 2% DSS feeding (Fig 5F). FMT partially restored the colon length of IEC^ΔChil1^ mice to that of Villin-cre mice after the DSS challenge, but did not have an impact on body weight loss or the level of inflammation in the gut (Fig. 5G-I). IEC^ΔChil1^ mice transplanted with *Lactobacillus* displayed a similar colitis phenotype as Villin-cre mice, characterized by similar weight loss, colon length reduction, and gut inflammation (Fig. 5G-I). These findings further validate the notion that Chi3l1-regulated gut microbiota, especially *Lactobacillus*, offers protection against colitis.

## Discussion

The intricate relationship between gut bacteria and their host organisms is crucial for health, maintained through co-evolution and co-speciation. Understanding how the host influences these bacterial communities is critical to understand the complex interplay between host and gut bacteria. In this study, we discovered that Chi3l1 interacts with gut microbiota via peptidoglycan, aiding the mucus colonization of beneficial bacteria like *Lactobacillus*, which protects against colitis (Fig. 6). <u>Our “scaffold model” demonstrates that Chi3l1 binds to bacterial peptidoglycan,</u> <u>helping anchor and organize bacteria within the mucus layer. This interaction promotes the</u> <u>colonization of beneficial bacteria, enhancing gut health.</u>

**Figure 6.**
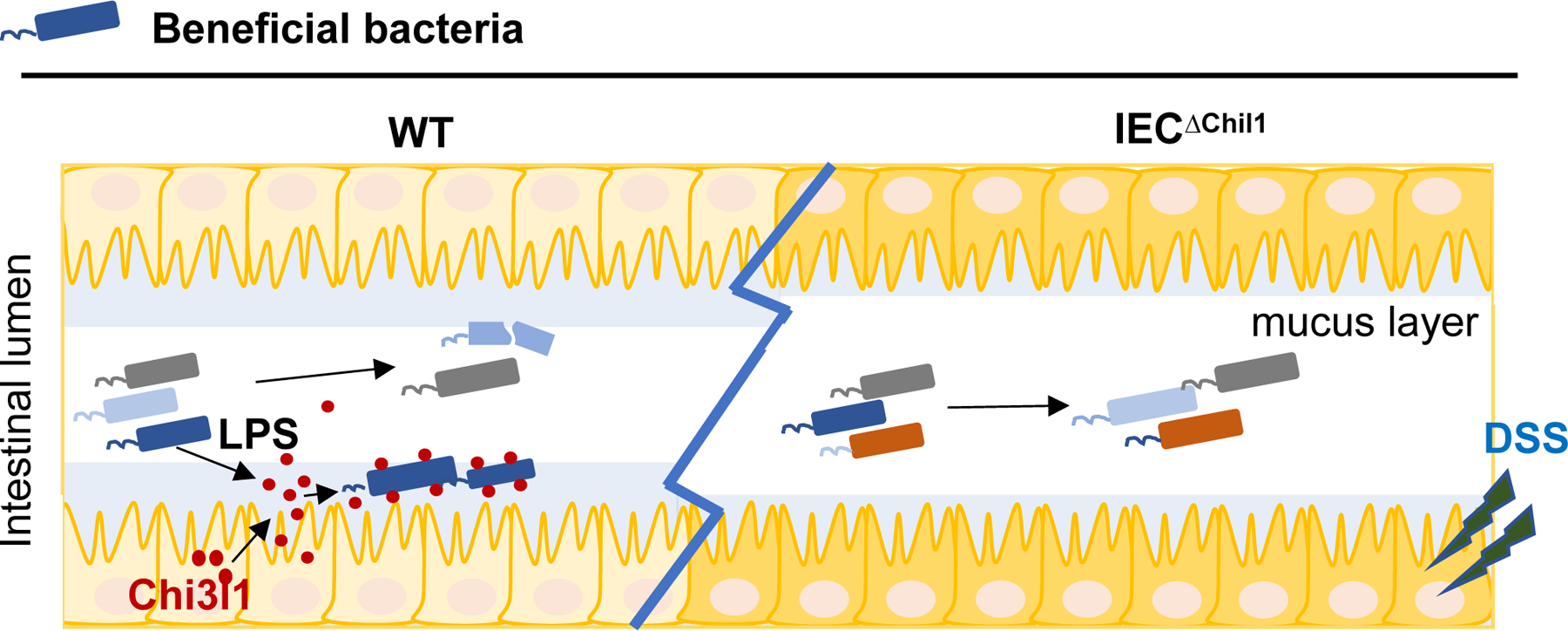
A schematic working model. Intestinal epithelial cells are stimulated by the gut microbiota to express Chi3l1. Once expressed, Chi3l1 is secreted into the mucus layer where it interacts with the gut microbiota, specifically through a component of bacterial cell walls called peptidoglycan. This interaction between Chi3l1 and bacteria is beneficial for the colonization of bacteria in the mucus, particularly for gram-positive bacteria like *Lactobacillus*. Moreover, a deficiency of Chi3l1 leads to an imbalance in the gut microbiota, which exacerbates colitis induced by dextran sodium sulfate (DSS).

Several studies have suggested a potential correlation between Chi3l1 and bacteria. For instance, it has been demonstrated that Chi3l1 is produced by intestinal epithelial cells in enteritis disease and aids in the elimination of pathogenic bacteria in the gut by neutrophils[24]. Another study observed an increase in Chi3l1 expression in mammary tissues of dairy cows infected with pathogenic *E. coli*, which promoted the recruitment of neutrophils[25]. Furthermore, Chi3l1 was found to be induced during *Streptococcus pneumoniae* infection, where it enhanced bacterial elimination by preventing the death of lung macrophages and improving host tolerance[26]. <u>One study demonstrated that Chi3l1 contributes to the pathogenesis of colitis, possibly by facilitating the adhesion and invasion of bacteria onto colonic epithelial cells. However, we observed the presence of the Gram-negative bacterium *Salmonella typhimurium* in the mice used in their study[27]. According to standard for SPF-grade mice, Salmonella should not be present in laboratory mice, and it was also not detected by our 16S RNA sequencing. However, if mice are inadvertently infected with Salmonella in the context of IBD, it could likely exacerbate the development of IBD. Our data also demonstrate that Chi3l1 can bind to Gram-negative bacteria (Figure 2B). Therefore, the role of Chi3l1 on microbiota or IBD is heavily dependent on the environmental context. Another study found that caffeine treatment reduce Chi3l1 RNA expression levels, alleviate colitis, and decrease bacterial translocation[28]. However, previous research suggested that caffeine can directly inhibit inflammasome activation by suppressing MAPK/NF-κB signaling pathways[29]. Therefore, caffeine treatment may directly reduce colitis by inhibiting inflammation. Moreover, an additional control group of mice with Chi3l1 specifically knocked out in the gut is needed to examine the role of caffeine in the alleviation of colitis. However, the detailed mechanism of the interaction between Chi3l1 and the gut microbiota remains incompletely understood, partly due to the absence of Chi3l1-specific knockout mice and variations in mouse husbandry conditions</u>. Here, by using intestinal Chi3l1-specific knockout mice, we demonstrated a new function of Chi3l1 in gut, which shapes bacterial colonozation through direct interaction with bacterial cell wall component, PGN. Moreover, Chi3l1-regulated gut microbiota, especially *Lactobacillus*, offers protection against colitis.

The bacterial cell wall is a complex structure mainly composed of peptidoglycan, but it may also contain other components such as teichoic acids and lipoteichoic acid in gram-positive bacteria, or an outer membrane containing various polysaccharides, lipids, and proteins in gram-negative bacteria[30]. Our findings indicated that lipopolysaccharide (LPS), a component of the bacterial cell wall, can slightly increase Chi3l1 expression (Fig. 1F&G), but higher levels of LPS do not further enhance Chi3l1 expression (data not shown). This suggests that there might be other components of the cell wall that can induce Chi3l1 expression in intestinal epithelial cells. Given the structural similarity between chitin and peptidoglycan (Fig. 2A), it is likely that Chi3l1 binds to the polysaccharide chains rather than the tetrapeptide in peptidoglycan. Previous studies have investigated the crystal structure of human Chi3l1 in complex with chitin, revealing a binding groove with different subsites for chitin fragments[31]. Other studies have also identified specific amino acids in chitinases that play a key role in the interaction with chitin[32, 33]. Nevertheless, further investigation is necessary to better understand the binding sites in Chi3l1 for peptidoglycan.

In the colon, there are two distinct parts of the mucus layer. The inner layer is attached and has a low number of microbes, while the outer layer is looser and densely populated by microorganisms. [34]. The diversity of bacteria in the mucus layer is similar to that found in the gut lumen[35]. It has to be noted that not all Gram-positive bacteria were reduced in Chil1^-/-^ mice, as we found an increase in *Turicibacter* in the colon (Fig. 3G) and feces (data not shown). We believe this may be due to a combined effect of the host and gut microbiota. Another important factor for microbial growth in the colon is the integrity of the mucus barrier[11].We noticed a thinning of the mucus barrier in the mice lacking Chi3l1 compared to normal mice (Supplementary Figure 3 A&B).

However, we did not observe an increase in mucin-degrading bacteria such as *Bacteroides* or *Allobaculum*[36, 37] or a decrease in mucin-producing cells in these mice. We think that there may be bacteria that aid in the formation of the gut mucus, and these bacteria are decreased in mice lacking Chi3l1. <u>Recent studies have introduced an “encapsulation model” regarding the nature of mucus in the colon. According to this model, the mucus in the proximal colon forms a primary encapsulation barrier around fecal material, while the mucus in the distal colon forms a secondary barrier.[38] Our findings indicate that Chi3l1 is expressed throughout the entire colon, including the proximal, middle, and distal sections (data not shown). This suggests that Chi3l1 likely promotes bacterial colonization across the entire colon. Despite most mucus being expelled with feces, the constant production of mucus and the minimal presence of Chi3l1 in feces (Figure 4C) indicate that Chi3l1 continuously plays a role in promoting the colonization of microbiota</u>.

In patients with inflammatory bowel disease (IBD), the density and diversity of the microbial community in the intestines are reduced. Specifically, there is a decrease in *Firmicutes* and an increase in *Bacteroides* and facultative anaerobic bacteria like *Enterobacteriaceae*[39]. However, the cause of colitis in IBD is still a subject of debate. Our data suggest that disruptions in the gut microbiota contribute to colitis. In summary, our study demonstrates that bacterial challenge induces the expression of Chi3l1 in intestinal epithelial cells. Once produced, Chi3l1 is released into the mucus layer where it interacts with the gut microbiota, particularly through peptidoglycan, a primary component of bacterial cell walls. This interaction is beneficial for the colonization of bacteria, especially gram-positive bacteria like *Lactobacillus*, in the mucus layer. Dysbiosis resulting from a lack of Chi3l1 exacerbates DSS-induced colitis, highlighting the role of dysbiosis as a contributing factor to colitis.

## Methods

### Animal experiments and procedures

C57BL/6J (Strain No. N000013), Germ-free (Strain No. N000295), Chil1^fl/fl^ (Strain No. T013652), Chil1^-/-^ (Strain No. T014402) mice were purchased from GemPharmatech. Villin-cre mice were provided by Dr. Qun Lu (Yunnan University, China,). All mouse colonies were maintained at the animal core facility of Yunnan University. C57BL/6J was used as wildtype control since Chil1^-/-^ mice are on the C57BL/6J background, as determined by PCR (data not shown). The animal studies described have been approved by the Yunnan University Institutional Animal Care and Use Committee (IACUC, Approval No. YNU20220256). Female mice aged 8-10 weeks old were used in most studies.

*Genotyping* <u>Tail clippings were placed in 1.5 ml tubes with 75 µL of master mix solution (60 µL H₂O, 7.5 µL 250 mM NaOH, 7.5 µL 2 mM EDTA) and incubated at 98°C for 1 hour. After cooling to 15°C, 75 µL of neutralization buffer (40 mM Tris HCl, pH 5.5) was added, and the samples were centrifuged at 4000 rpm for 3 minutes. A 1:10 dilution was prepared by mixing 2 µL of supernatant with 18 µL of water for genotyping PCR. A 25 µL PCR reaction mix was prepared with 12.5 µL of 2x Taq Master Mix (Dye Plus, Vazyme P112-03), 1 µL of each primer, 2 µL of template, and water. PCR was conducted using a Biorad machine with the following program: 95°C for 5 minutes; 98°C for 30 seconds; 65°C with a 0.5°C decrement per cycle for 30 seconds; 72°C for 45 seconds, repeating steps 2-4 for 20 cycles; 98°C for 30 seconds; 55°C for 30 seconds; 72°C for 45 seconds, repeating steps 5-7 for 20 cycles; 72°C for 5 minutes; and hold at 10°C. Primer sequences are listed in Table S1</u>.

**Table S1.**
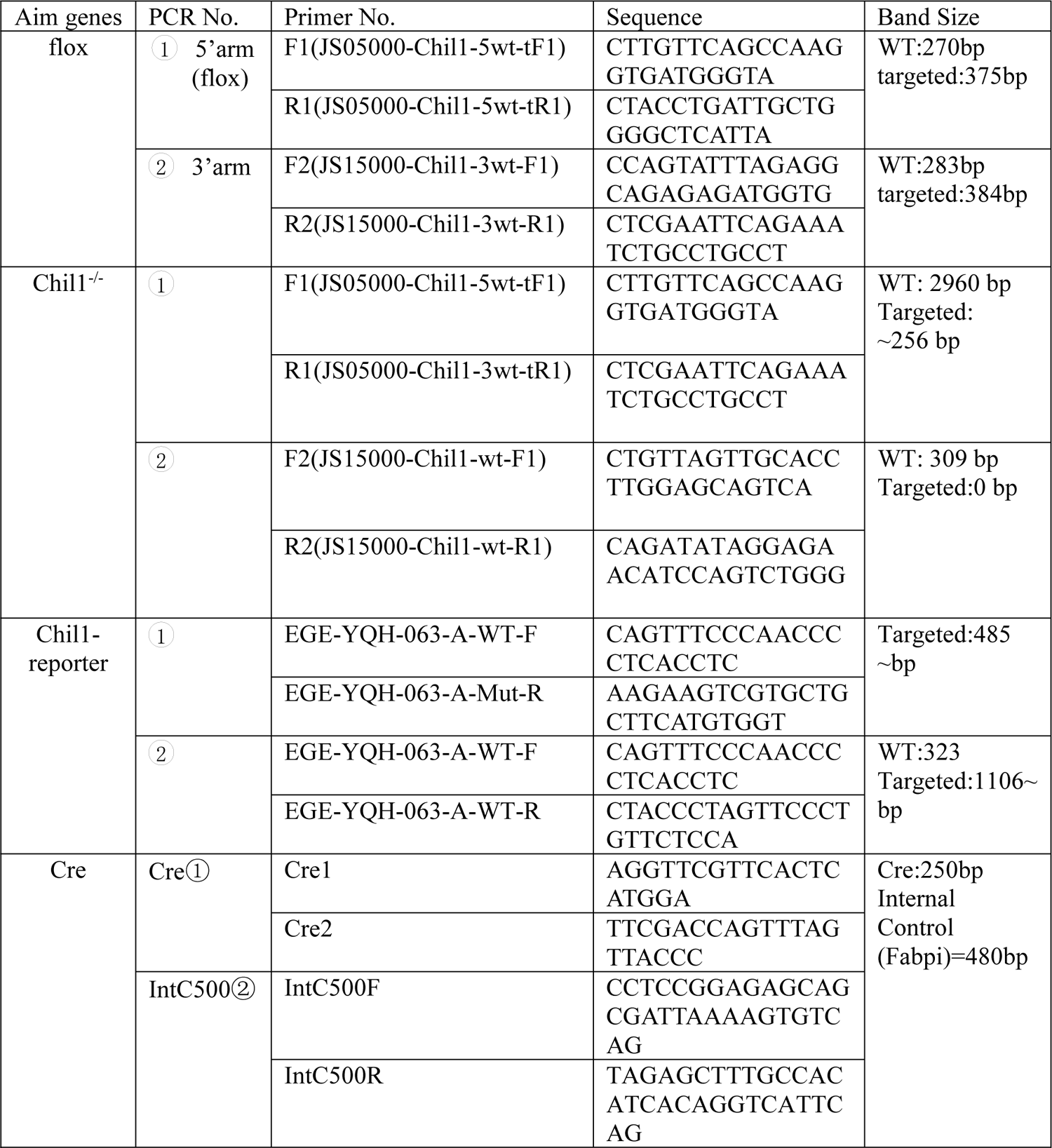

*Rectal administration of FDAA-labeled E. faecalis* For FDAA-labeled *E. faecalis*, *E. faecalis* were grown in LB media until reaching mid-exponential phase, approximately 4 hours. FDAA was then added to the culture media to a final concentration of about 17 µM. The *E. faecalis* continued to grow for 4 hours and were then harvested by centrifugation (5000 x g for 10 min), washed, and resuspended in PBS at a density of 5x10^9^ CFU/ml. For rectal administration, wildtype and Chil1^-/-^ mice were fasted overnight (5:00pm to 9:00am) before intraperitoneally (i.p.) injecting them with 400mg/kg tribromoethanol (Nanjing AIBI BioTechnology, M2910), followed by rectal injection of 1x10^9^ FDAA-labeled E. faecalis in 200 μl PBS via a flexible catheter. The catheter was inserted into the anus to a depth of 4.5 cm, and the *E. faecalis* were injected slowly to avoid overflow. The mice were then kept upside down for approximately 2 minutes. After 4 hours of rectal injection, the mice’s colons were collected and immediately embedded in OTC embedding medium for observation on 5μm thick frozen sections. OP50-mCherry (1x10^9^ <u>CFU/mice) treated same as *E. faecalis*.</u>

*DSS-induced mouse colitis* Villin-cre and IEC^ΔChil1^ mice were fed a 2% dextran sodium sulfate (DSS, MP, 9011-18-1) solution in their drinking water for 7 days. The mice’s body weight was monitored daily during the feeding period. After the 7-day DSS treatment, the mice were sacrificed. Colons were collected, and their length was measured from the cecum to the rectum. Colon paraffin sections were harvested, and H&E staining was performed to examine gut inflammation.

*Antibiotics treatment* IEC^ΔChil1^ mice were fed an antibiotics mixture containing 0.5 mg/ml of Metronidazole (Solarbio, M8060), 1 mg/ml of vancomycin (Solarbio, V8050), 1 mg/ml of Ampicillin (Solarbio, A8180), and 0.5 mg/ml of Neomycin Sulfate (Solarbio, N8090) in their drinking water for 7 days (approximately 5ml per mouse per day). After the 7-day antibiotics treatment, the mice were orally gavaged with 200μl of the antibiotics mixture for another 3 days. Microbiota depletion was examined in feces using 16S rRNA qPCR on the 10th day of antibiotics feeding.

*Fecal microbiota transplantation (FMT)* Fresh feces were collected from 8-10 weeks old Villin-cre mice and immediately snap-frozen in liquid nitrogen. On the experimental day, feces were <u>resuspended</u> in PBS to a concentration of 200 mg/ml and centrifuged at 350g for 5 minutes to collect the supernatant. Antibiotics pre-treated IEC^ΔChil1^ mice were orally gavaged with 10 μl of the dissolved feces per gram of mouse weight for 14 days.

*Oral gavage of Lactobacillus reuteri Lactobacillus reuteri* was grown in MRS broth at 37°C for 48 hours under anaerobic conditions. The bacteria were harvested by centrifugation (5000 x g for 10 min), washed, and resuspended in PBS at a density of OD_605_=1.2-1.3/ml. Antibiotics pre-treated IEC^ΔChil1^ ^mice^ were orally gavaged with 200μl of the dissolved Lactobacillus reuteri per mouse for 14 days.

### Bacterial colony identification

Fresh feces were collected from 8-week-old wild-type mice and dissolved in LB culture medium. The dissolved feces were then cultured at 37°C for 12 hours and plated onto LB agar plates. The grown colonies were picked using sterile pipette tips and resuspended in 20 μl of sterile water. A PCR reaction was performed using 2 μl of the bacterial suspension as template DNA and universal bacterial 16S rRNA primers (27F, 5’-AGAGTTTGATCCTGGCTCAG-3’ and 1492R, 5’-GGTTACCTTGTTACGACTT-3’) with reaction conditions: 95°C for 5 min followed by 35 cycles of 95°C for 30sec, 55°C for 30sec, 72°C for 2min and then 72°C for 20min. The amplicons were then sequenced, and the resulting sequences were analyzed using BLASTN and the NCBI database for species identification.

### Treatment of DLD-1 cells with live, heat-killed bacteria or LPS

Bacteria harvested from mouse feces or specific strains were grown in LB medium at 37°C for 12 hours under aerobic conditions. The bacteria were then collected by centrifugation (5000 x g for 10 min), washed, and resuspended in PBS at a density of 1.2 x 10^10^ CFU/ml. Live bacteria were used directly, while heat-killed bacteria required further heating at 80°C for 30 minutes. DLD-1 cells were grown in DMEM medium supplemented with 10% FBS. Prior to treatment, the cells were replated and allowed to reach 80% confluency. Either live or heat-killed bacteria were added to the cells at a multiplicity of infection of 20 each well, and 100 pg/ml LPS treatment(Sigma, L4392; diluted in PBS) was also performed. After 12-hour incubation, cellular proteins were extracted for western blot analysis or the cells were subjected to immunofluorescent staining.

### Cell Culture and Bacterial Adhesion Assay

<u>DLD-1 cells were cultured in 6-well plates until they reached approximately 80% confluency per well. The cells were then washed three times with PBS to remove any residual antibiotics. Fresh DMEM supplemented with 10% FBS (without antibiotics) was added to the cells. E. coli strains were introduced to each well at a multiplicity of infection (MOI) of 20 and incubated for 2.5 hours at 37°C. After incubation, the cells were washed three times with PBS. For the adhesion assays, the cells were lysed with 1 mL of 1% Triton-X100 (BBI, 9002-93-1) in deionized water for 30 minutes. The cell lysates were then plated onto Luria broth (LB) agar plates at various dilutions and incubated overnight at 37°C. The following day, colonies were counted and calculated as colony-forming units (CFU)/mL</u>.

### shRNA Lentivirus Packaging and Transfection

<u>For the packaging of shRNA lentivirus, shRNA control and Chi3l1 shRNA1 lentiviruses were packaged in 293T cells. The cell culture medium containing the virus was collected and stored at - 80°C. For transfection of DLD-1 cells, the cells were cultured in 6-well plates until they reached approximately 80% confluency. The cell culture medium was then replaced with a mixture of culture medium containing the virus and fresh cell culture medium in a 1:1 ratio. Following transfection, the bacterial adhesion assay was performed as described above</u>.

### Bacterial or peptidoglycan binding assay

Different bacteria strains were grown in LB medium at 37°C for 12 hours under aerobic conditions. The bacteria were collected by centrifugation (5000 x g for 10 min), washed, and resuspended in MES buffer (25mM MES, 25mM NaCl, Ph=6.0) at a density of 5x10^9^ CFU/ml. 1μg Recombinant mouse Chi3l1 (rmChi3l1) was added to the bacterial suspension and incubated at 4°C under rotation overnight. Supernatant, wash fractions, and bacterial-bound fractions were collected and analyzed using western blot analysis. For the peptidoglycan binding assay, 1μg rmChi3l1 or recombinant human Chi3l1 (rhChi3l1) and bovine serum albumin (BSA) was incubated with 100μg peptidoglycan (PGN). The incubation and wash procedure were similar to the bacterial binding assay, and the proteins in each fraction were analyzed using silver staining or western blot.

### Hematoxylin/eosin(H&E) and Periodic acid-Schiff and Alcian blue (AB-PAS) staining

Tissues were fixed with buffered 10% paraformaldehyde (BI, E672001-0500) overnight at 4°C and embedded in paraffin. Ultra-thin tissue slices (5μm) were prepared and deparaffinized. H&E staining was performed on the tissue sections, and slides were examined under a microscope (Leica). For AB-PAS staining, tissues were fixed in Carnoy’s solution (60% Ethanol, 30% Chloroform, 10% Acetic Acid) for 24 hour at 4°C and embedded in paraffin. AB-PAS staining (Solarbio, G1285) was performed according to the manufacturer’s protocol. The staining was visualized under the microscope (Leica).

### Immunohistochemical (IHC) and immunofluorescent (IF) staining

Tissue paraffin sections were prepared as previously described for H&E staining. Antigen retrieval was performed by treating the sections with citric acid (pH 6.0) at 95°C for 15 minutes, followed by cooling to room temperature. The sections were then washed with PBS and ddH2O. To block any nonspecific binding, a blocking buffer containing 5% goat serum, 3% BSA, and 0.1% Triton X-100 in PBS was applied to the sections for 1 hour at room temperature in a humidity chamber. The sections were then incubated with anti-Chi3l1 primary antibodies (Invitrogen, PA5-95897, 1:200), followed by staining with Goat anti-Rabbit IgG (H+L) Secondary Antibody-Biotin (Invitrogen, 65-614, 1:2000). Finally, the sections were stained with Horseradish Peroxidase conjugate antibody (Invitrogen, A2664, 1:2000) and developed with DAB for 10 minutes. The slides were examined under a microscope (Leica).

Immunofluorescent staining was also performed on paraffin sections using specific primary antibodies. These included anti-MUC2 (Invitrogen, PA5-103083, 1:50), anti-Chi3l1 (Abcam, ab180569, 1:400; antigen retrieval with Tris-EDTA at pH 9.0), anti-ChgA (Santa Cruz, sc-393941, 1:200; antigen retrieval with citrate at pH 6.0), anti-UEA-1-FITC (GeneTeX, GTX0151, 1:200; antigen retrieval with citrate at pH 6.0), and anti-LTA (Invitrogen, MA1-7402, 1:50). The secondary antibodies used were 488-conjugated Affinipure Goat Anti-Rabbit IgG(H+L) (Jackson ImmunoResearch, 111-545-003, 1:1000) and 594-conjugated Goat Anti-Mouse IgG (H+L) (Jackson ImmunoResearch, 115-585-003, 1:1000). Slides were washed and mounted with antifade medium (Vectashield, H-1000-10). Nuclei were stained with DAPI (Beyotime, c1006, prediluted). Images were captured using a fluorescence microscope (Leica, 2084 DP-80).

Immunofluorescent staining on DLD-1 cells, cells were seeded on coverslips in 12-well plate and challenged with 100 pg/ml LPS (Sigma, L4391; diluted in PBS) for 12 hours. Cells were washed with cold-PBS twice gently, then fixed with 2% paraformaldehyde in PBS at room temperature for 10 min. After removal of fixation buffer and wash twice with cold-PBS, cells were blocked with blocking buffer (3% BSA, 0.5% Triton-X-100 in PBS) for 1 hour in humidity chamber at room temperature. Rabbit anti-Chi3l1 antibody (Proteintech, 12036-1-AP; 1:200) was applied at room temperature for 1 hour. After wash with 1 x TBST three times, secondary antibody AlexaFluor 488 (Jackson ImmunoResearch, 111-545-003, 1:1000) was applied for another 1 hour at room temperature in a humidified chamber under darkness. Finally, cells were counterstained with DAPI (beyotime, C1006) and mounted onto slides. Images were captured using an Olympus BX53F2 microscope.

### Fluorescence in situ hybridization (FISH)

Murine intestinal paraffin sections were prepared according to the previously described method for H&E staining. The tissues sections were rehydrated using a graded ethanol series and then washed with distilled water. The gram-positive bacterial probe, consisting of three different sequences (/5Alex550N/TGGAAGATTCCCTACTGC/3AlexF550N/, /5Alex550N/CGGAAGATTCCCTACTGC/3AlexF550N/, /5Alex550N/CCGAAGATTCCCTACTGC/3AlexF550N/), or the control nonspecific probe (/5Alex550N/ACTCCTACGGGAGGCAGC/3AlexF550N/), was diluted to a concentration of 100 nM in FISH hybridization buffer (containing 20 mM Tris pH 7.2, 0.9 M NaCl, and 0.1% SDS) and applied to the slides. The slides were then incubated overnight at 56°C in a humidified chamber. Following incubation, the slides were washed and the nuclei were counterstained with DAPI. The images were captured using a fluorescence microscope (Leica, 2084 DP-80).

### Immunoblot and silver staining

Protein extraction from cultured cells involved lysing the cells in 2% SDS lysis buffer, which is prepared by dissolving 2g of SDS powder in 100ml of sterilized ddH2O. Bacteria or peptidoglycan precipitates were resuspended in MES buffer, which contained 25mM MES, 25mM NaCl, and had a pH of 6.0. For protein extraction from mice ileum and colon tissues, 30mg of snap-frozen tissues were homogenized in 1ml of RIPA buffer, which contained 10mM Tris-HCl (pH 8.0), 1mM EDTA, 0.5mM EGTA, 1% TritonX-100, 0.1% sodium deoxycholate, 0.1% SDS, 140mM NaCl, 1mM PMSF, and a proteinase inhibitor. The lysates were then supplemented with 5x SDS loading buffer to a final concentration of 1x. The resulting mixture was subsequently boiled at 100°C for 10 minutes and centrifuged at 4°C and 12,000 rpm for 10 minutes. The supernatants were collected for western blot analysis. The supernatants were separated using a 10% SDS-PAGE gel and then transferred to a polyvinylidene fluoride membrane. The membranes were blocked with 5% nonfat milk in TBST buffer (containing 0.1% Tween-20 in tris-buffered saline) and sequentially incubated with primary antibodies and appropriate horseradish peroxidase (HRP)-conjugated secondary antibodies. Protein bands were detected using enhanced chemiluminescence (ECL) reagent with a Minichemi Chemiluminescence Imaging System. The primary antibodies used included anti-Chi3l1 (RD, MAB2649, 1:2000) and anti-alpha-Actinin (Cell Signaling, 69758S, 1:1000). The secondary antibodies used were goat anti-Rat-IgG (Cell Signaling, 7077s, 1:1000) and goat anti-mouse (Invitrogen, 62-6520, 1:1000). For silver staining, the PAGE gel was subjected to silver staining using a fast-silver staining kit (Beyotime, P0017S) following the manufacturer’s instructions.

### DNA extraction for 16S rRNA analysis

For isolation of luminal contents from murine ileum and colon, a 9 cm section of ileum and 3 cm section of colon were cut open longitudinally, and luminal contents were scratched off into a pre-weighed 2-ml sterile freezing vial. The weight of the contents was recorded for further processing. For isolation of mucus from murine ileum and colon, tissues were flushed with 2 ml of ice-cold PBS into a pre-weighed 2-ml sterile freezing vial after removal of the liminal contents. The mucus was then pelleted by centrifugation at 10,000g for 10 minutes, and the supernatant was removed. For feces isolation, fresh murine feces were collected into a sterile freezing vial weighing 2 ml, and immediately snap-frozen in liquid nitrogen. For bacterial DNA extraction, DNA was extracted and purified following the manufacture’s protocol using the EIANamp stool DNA kit (TIANGEN, DP328-02).

### Microbiota 16S rRNA gene sequencing

Fecal samples, ileum contents, and colon contents were collected from wildtype, Chil1^-/-^ littermates, or Villin-cre, IEC^ΔChil1^ littermates and immediately frozen in liquid nitrogen. The microbial genomic DNA was extracted and 16S rRNA sequencing was performed by Biomarker Technologies. The hypervariable regions V3 and V4 of the bacterial 16S rRNA gene were sequenced using universal primers that flank these regions V3 (338F 5’-ACTCCTACGGGAGGCAGCA-3’) and V4 (806R 5’-GGACTACHVGGGTWTCTAAT-3’). The sequencing was done using the IIIumina Sequencing platform. The resulting 16S rRNA gene sequences were analyzed using scripts from the BMK Cloud platform (www.biocloud.net). The microbial classification was performed using the SILVA138 and hierarchical clustering algorithms. The OTUs were determined by clustering the sequences with 97% similarity and were classified into different taxonomic ranks. The relative abundance of each bacterial species was visualized using R software. The raw 16S rRNA gene sequencing data can be accessed on the BMKCloud platform under the project ID Microbial_updateReport_20211221092022313.

### 16S qPCR analysis

Quantitative PCR was performed using SYBR green master mix (Thermo Fisher, A25742) in triplicates. This was done on a Real-Time PCR QuatStudio1 with accompanying software, following the instructions provided by the manufacturer (Life Technologies, Grand Island, NY, USA). The abundance of specific bacterial groups in the intestine was determined using qPCR with either universal or bacteria-specific 16S rRNA gene primers. Standard curves were constructed with *E.Coli OP50* 16S rRNA gene, which was amplified using conserved 16S rRNA primers. It should be noted that qPCR measures the number of 16S gene copies per sample, not the actual bacterial numbers or colony forming units. Primer sequences are provided in table S2.

**Table S2.**
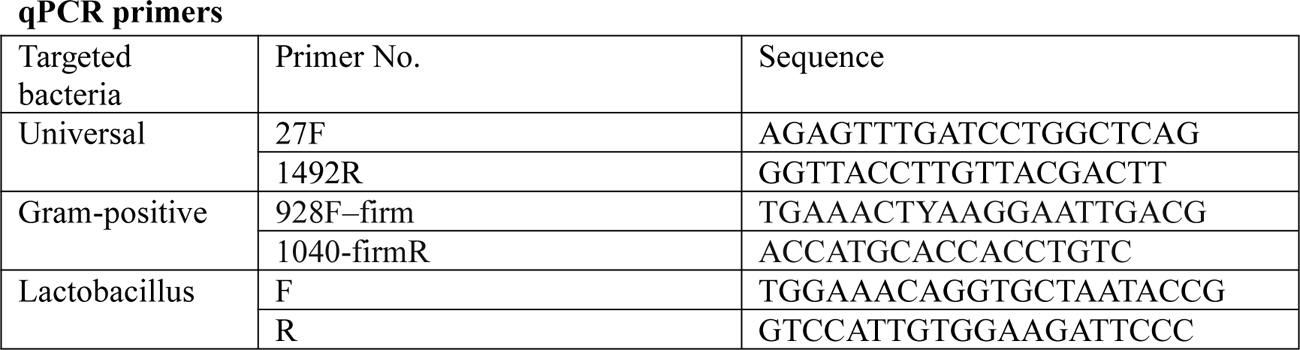

### Key source tables

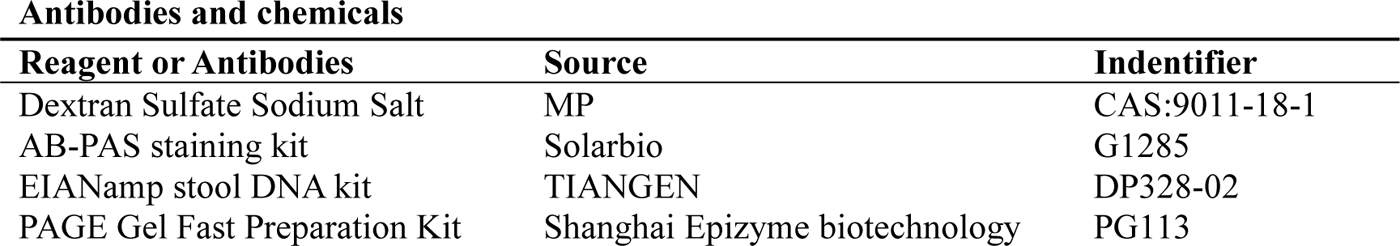
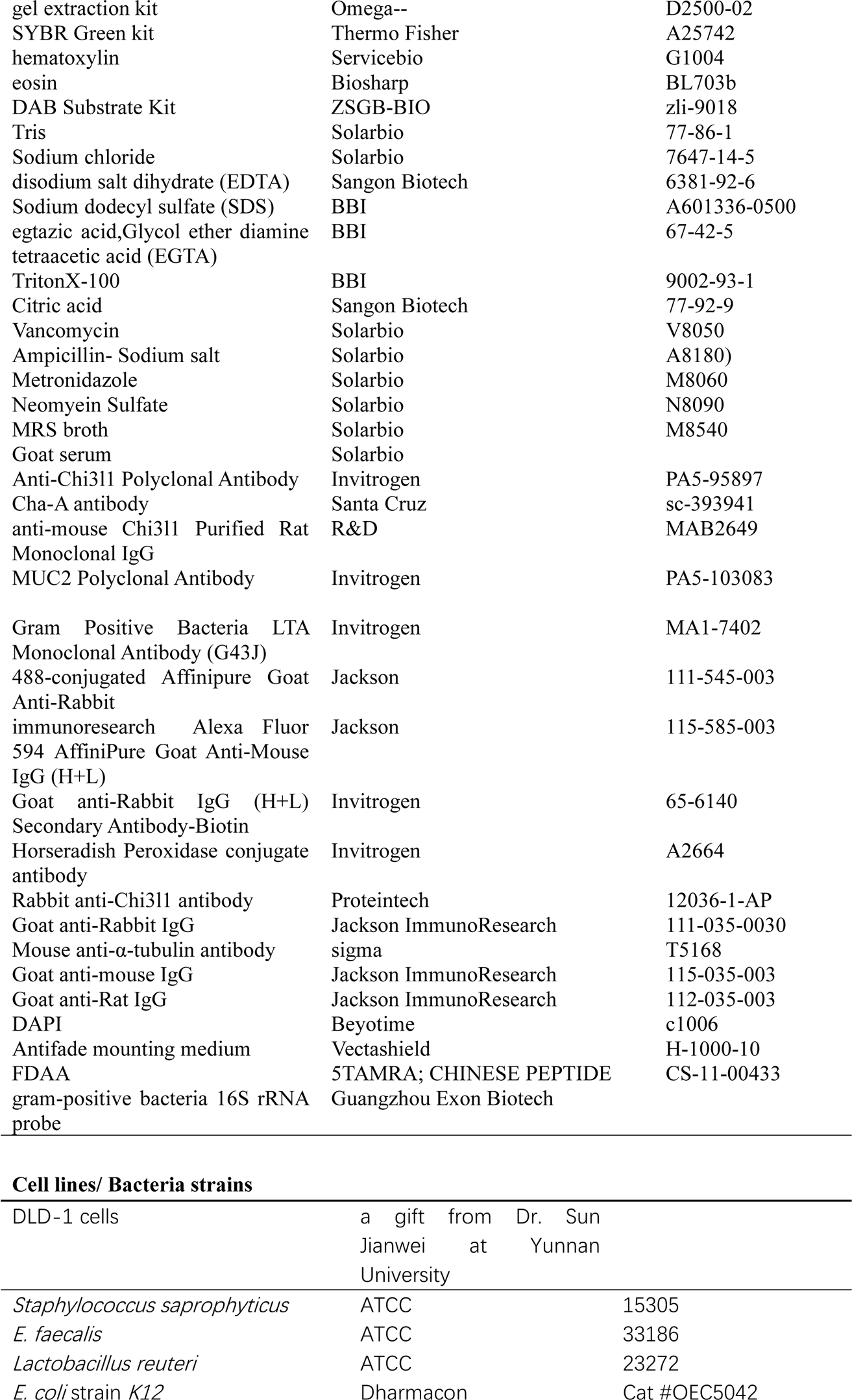
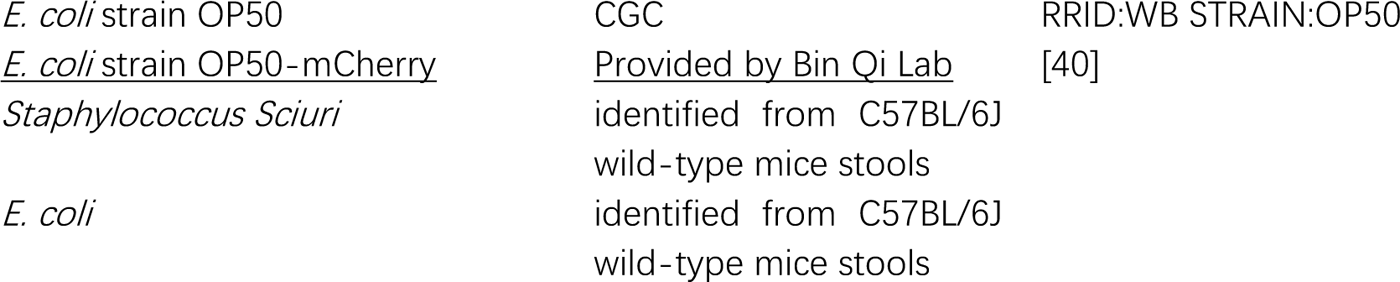

### Statistical analysis

Data were presented as mean ± SEM. Statistical analyses were carried out using GraphPad Prism (GraphPad Software). Comparisons between two groups were carried out using unpaired Student t test. Comparisons among multiple groups (n>=3) were carried out using one-way ANOVA. P values are as labeled and less than 0.05 was considered not significant. Patient data were analyzed by Mann-Whitney test.

## Author Contributions

Y.C. and R.Z.Y. performed experiments, analyzed data, and wrote the manuscript. B.Q. organized and designed the study. Z.S. initiated, organized, designed the study and wrote the manuscript.

## Acknowledgements

We thank Qun Lu (Yunnan University) for providing Villin-cre mice. We thank Wenxiang Fu (Yunnan University) for imaging technical support. We thank Jianwei Sun (Yunnan University) for providing DLD-1 cell line. We thank Zehan Hu (Shanghai Jiao Tong University) for methods of bacteria 16S FISH staining. This work was supported by Ministry of Science and Technology of China (2019YFA0803100, 2019YFA0802100 to B.Q.), National Natural Science Foundation of China (32071129 to Z.S., 32170794 to B.Q.), Yunnan Fundamental Research Projects (202101AT070022, 202001AW070006 to Z.S., 202201AT070196 to B.Q.), Science and Technological Talent Cultivation Plan of Yunnan Province (C619300A086 to Z.S., K264202230211 to B.Q.), Yunnan Provincial Science and Technology Project at Southwest United Graduate School (202302AP370005 to B.Q.)

