## Supplementary Figure 1-4 for "Peptidoglycan-Chi3l1 interaction shapes gut microbiota in intestinal mucus layer"

### Sup Figure 1

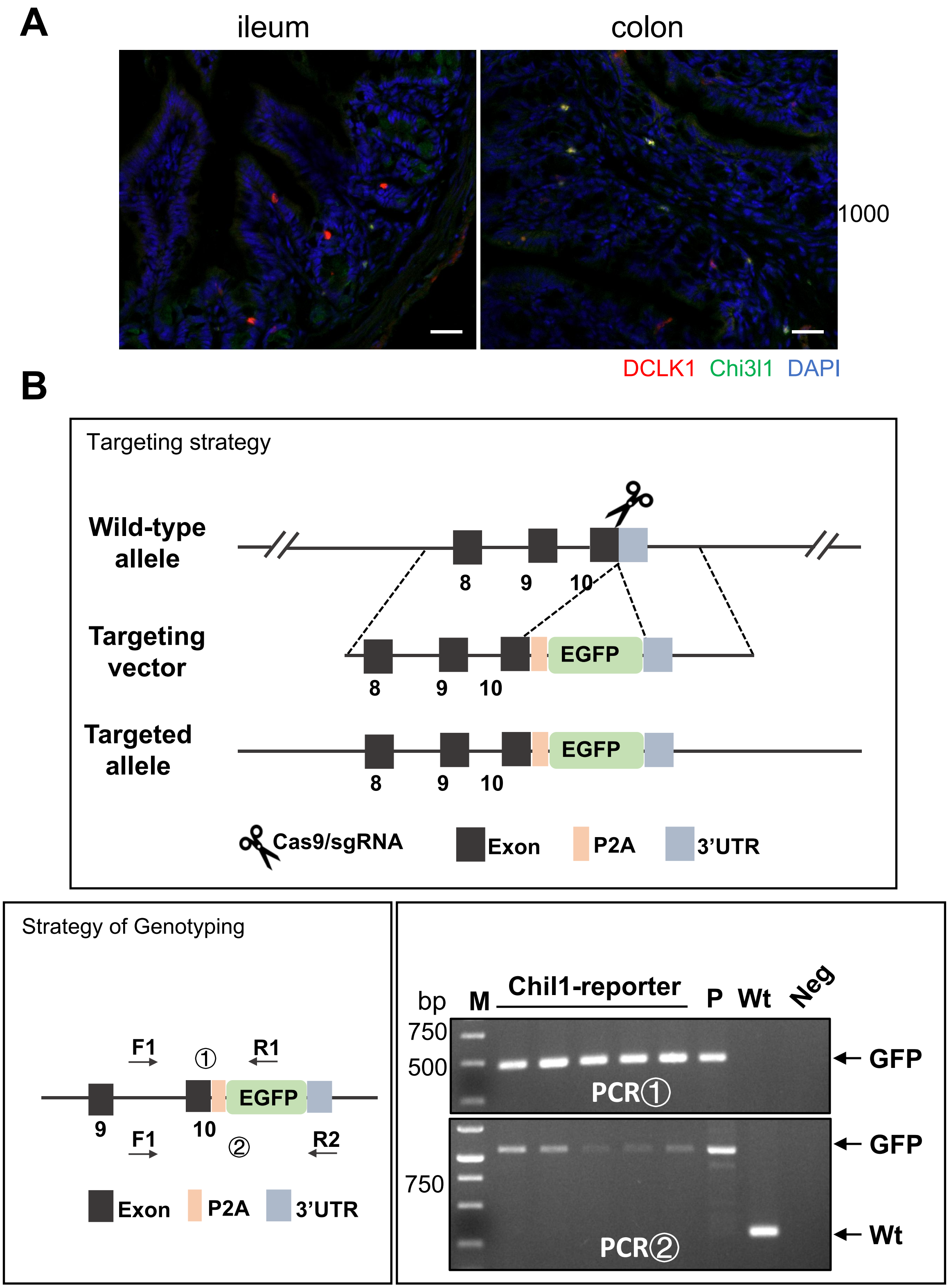
A
ileum
colon
1000
DCLK1 Chi3l1 DAPI
B
Targeting strategy
Wild-type allele
8
9
10
Targeting vector
8
9
10
EGFP
Targeted allele
8
9
10
EGFP
Cas9/sgRNA
Exon
P2A
3'UTR
Strategy of Genotyping
F1
①
R1
EGFP
9
10
F1
②
R2
Exon
P2A
3'UTR
bp
M
Chil1-reporter
P
Wt
Neg
750
500
GFP
PCR①
GFP
750
Wt
PCR②

**Supplementary Fig. 1 Chi3l1 don’t express in tuft cells.**
(A) Ileum and colon were collected from Chil1-EGFP reporter mice and stained with DCLK1 (red), Chi3l1(green), and nuclear DAPI (blue). Scale bar, 20μm. Representative images are shown, n=3 mice/group. (B) The construction, genotyping strategy, and genotyping results of Chil1-EGFP reporter mice. P: positive control; Wt: Wild-type control; Neg: Blank control(ddH2O); M: DNA Ladder.

Sup Figure 2

A

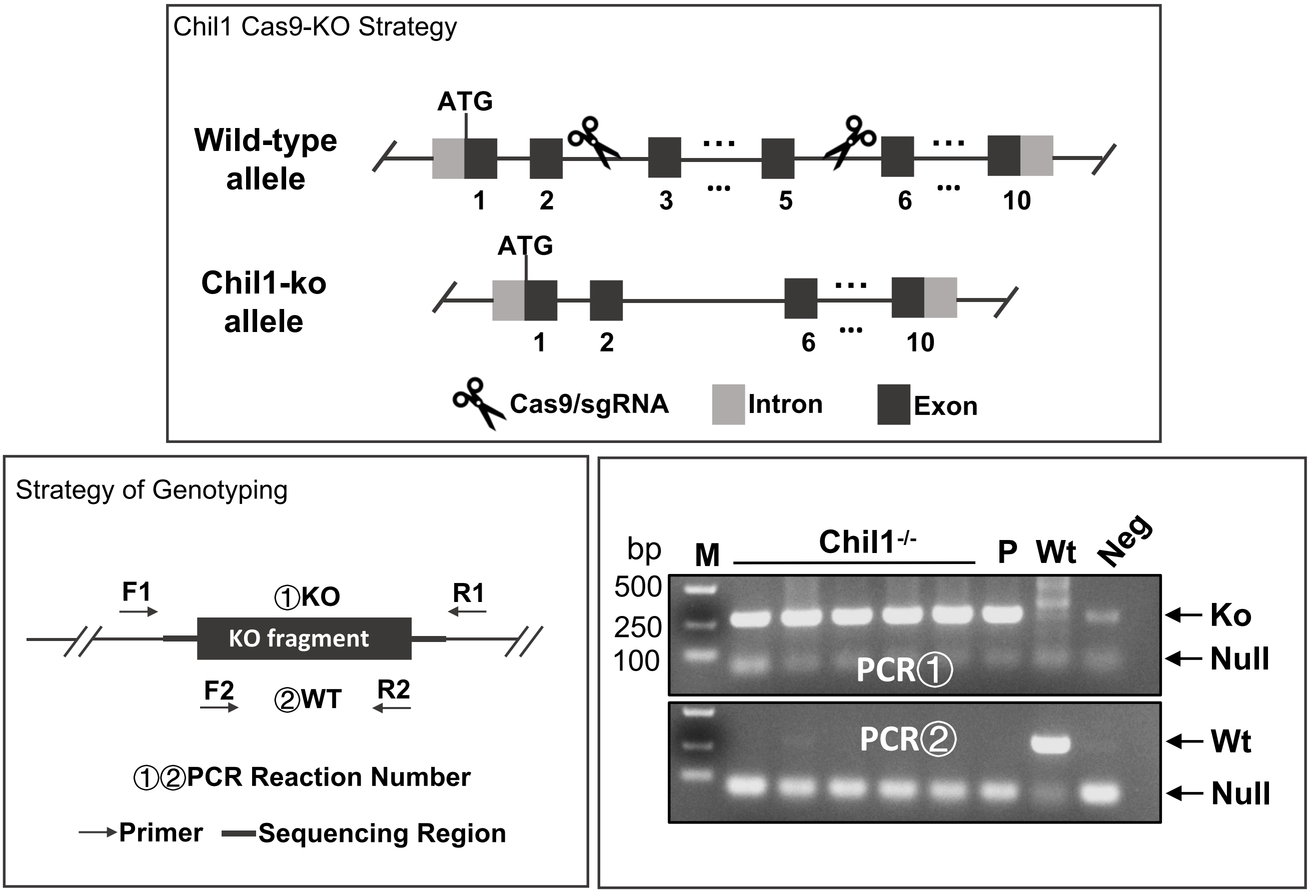


B

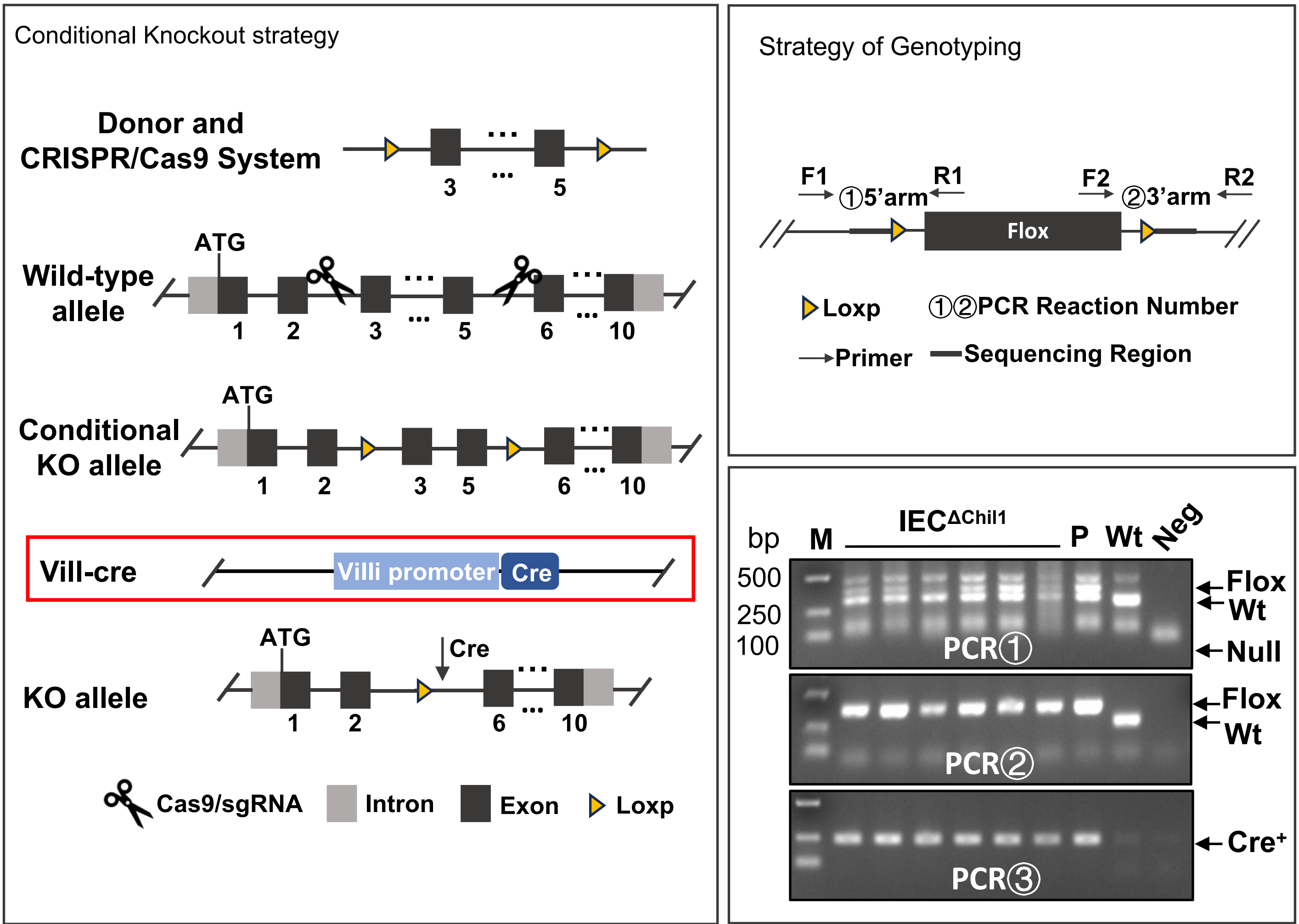

**Supplementary Fig. 2 The construction and genotype of Chi3l1$^{-/-}$ and IEC$^{\Delta Chil1}$ mice.**

(A) The construction, genotyping strategy and genotyping results of Chil1$^{-/-}$ mice. P: positive control; Wt: Wild-type control; Neg: Blank control(ddH2O); M: DNA Ladder.

(B) The construction, genotyping strategy and genotyping results of IEC$^{\Delta Chil1}$ mice. P:positive control; Wt: Wild-type control; Neg: Blank control(ddH2O); M: DNA Ladder. PCR① and ② implicated flox, ③implicated cre.

### Sup Figure 3

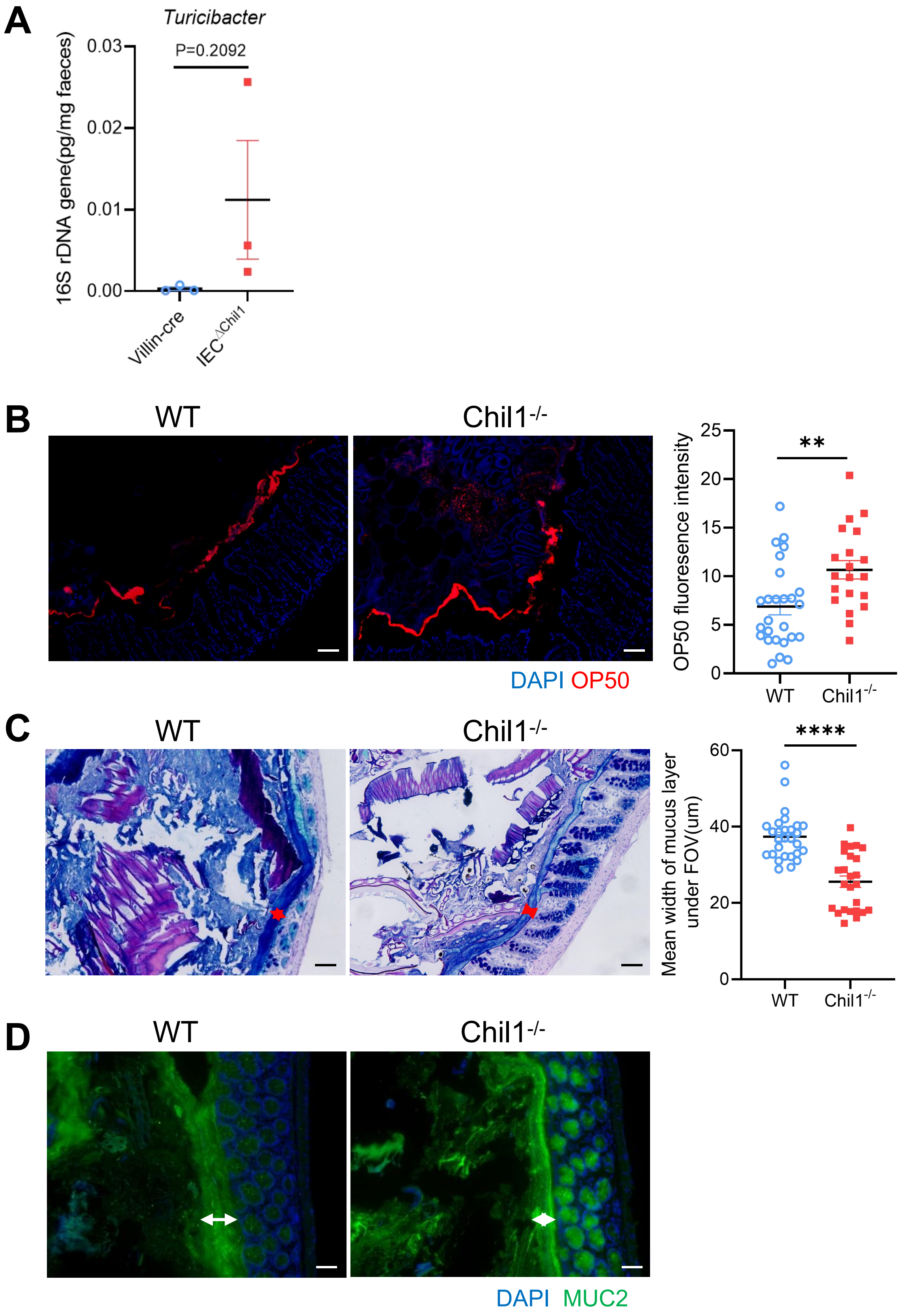

A
Turicibacter
16S rDNA gene(pg/mg faeces)
0.03
0.02
0.01
0.00
P=0.2092
Villin-cre
IEC^ΔChil1
B
WT
Chil1-/-
DAPI OP50
OP50 fluoresence intensity
25
20
15
10
5
0
**
WT
Chil1-/-
C
WT
Chil1-/-
Mean width of mucus layer
under FOV(um)
60
40
20
0
****
WT
Chil1-/-
D
WT
Chil1-/-
DAPI MUC2

**Supplementary Fig. 3 Chil1$^{-/-}$ mice possess shortening mucus layer.**
**(A)** qPCR analysis of *Turicibacter* in the feces of Villin-cre and IEC$^{\Delta Chil1}$ is shown. Values for each bacterial group are expressed relative to total 16S rRNA levels. **(B)** Rectal injection of both wildtype and Chi11$^{-/-}$ mice with mCherry-OP50 (a strain of E.coli expressing mCherry) for 4 hour. Colon sections were collected and colonization of OP50 was examined under microscope. Nuclei were stained with DAPI. n=3-4 mice/group.The average fluorescence intensity in each field of view (FOV) was analyzed. **(C)** Periodic acid-Schiff and Alcian blue (AB-PAS) staining in the colons of WT and Chil1$^{-/-}$ littermates. Scale bars, 100μm. The mean width of mucus layer in each field of view (FOV) was analyzed. **(D)** Immunofluorescence staining to detect Mucin 2 (green) and nuclear DAPI (blue) in colon from WT and Chil1$^{-/-}$ littermates. Scale bar, 50μm. Representative images are shown in C, D, n=4 mice/group.

### Sup Figure 4

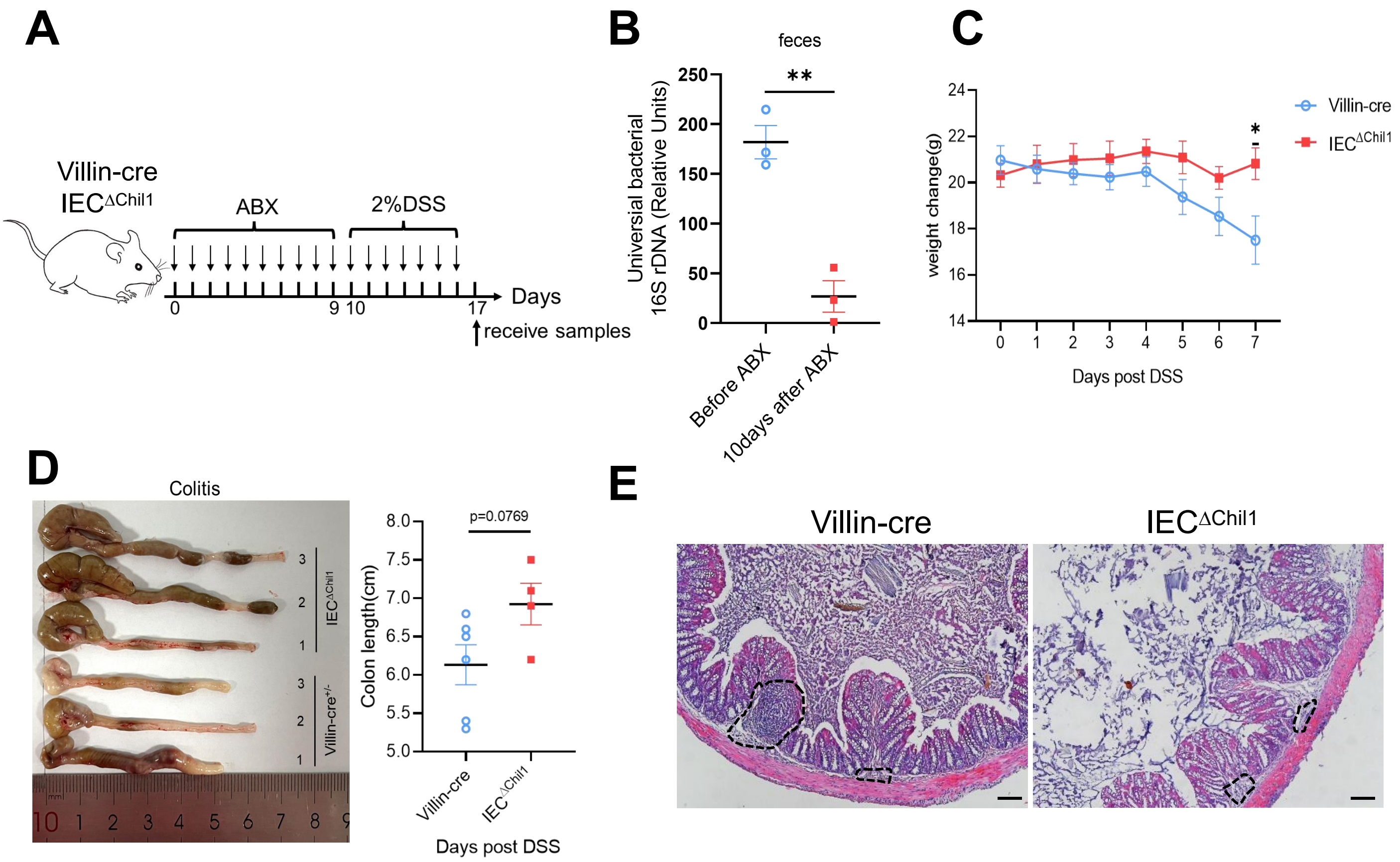


**Supplementary Fig. 4 Chi3l1-mediated bacteria, but not Chi3l1 itself affect more upon the development of colitis.**

**(A)** Schematic model of the experimental design. Both Villin-cre and IEC$^{\Delta Chil1}$ littermates were fed with 2% DSS in drinking water to induce colitis after elimination of gut microbiota by antibiotics for 10 days. **(B)** qPCR analysis of total bacteria in the feces of Villin-cre and IEC$^{\Delta Chil1}$ littermates. Values for each bacterial group are expressed relative to total 16S rRNA levels. Mean ± SEM is displayed. Two-tailed, unpaired student t-test was performed. **P<0.01, n=3/group. **(C)** Weight change of Villin-cre and IEC$^{\Delta Chil1}$ mice during DSS feeding. **(D)** Representative colonic length from Colitis Villin-cre and IEC$^{\Delta Chil1}$ mice (left) and the statistics of colonic length (right). Mean ± SEM is displayed. Two-tailed, unpaired student t-test was performed. P value is as indicated. **(E)** H&E staining of colitis mice colon from Villin-cre and IEC$^{\Delta Chil1}$. The inflamed area is outlined by black dotted lines, Scale bars=100 μm. n=4-6 mice/group.
